# On-The-Fly Live-Cell Intrinsic Morphological Drug and Genetic Screens by Gigapixel-per-second Spinning Arrayed Disk Imaging

**DOI:** 10.1101/2025.10.21.683625

**Authors:** Dickson M. D. Siu, Victor M. L. Wong, Bei Wang, Justin Wong, Alan S. L. Wong, Kevin K. Tsia

## Abstract

Next-generation drug discovery and functional genomics require rapid, unbiased single-cell profiling at scale—demands that exceed the limited speed, throughput, and labor-intensive labeling constraints of conventional high-content image-based screening. We introduce spinning arrayed disk (SpAD), a high-throughput, label-free imaging platform for live-cell imaging that integrates continuous circular scanning, ultrafast quantitative phase imaging (QPI), and a novel circular array of 96 culture chambers. SpAD achieves an order-of-magnitude reduction in imaging time compared to traditional fluorescence-based workflows, while remaining compatible with standard cell culture workflows. By extracting rich biophysical features using intrinsic morphological (InMorph) profiling and machine learning, SpAD enables sensitive, large-scale screening of drug responses and CRISPR gene knockouts without labeling. Critically, label-free biophysical readouts from SpAD reveal mechanism-linked changes in mass, refractive index, subcellular textures, and light scattering that fluorescent labels often obscure. SpAD thereby resolves subtle phenotypes and heterogeneous subpopulations with high reproducibility, providing a robust, scalable foundation for precision cellular morphological assays.

## Introduction

The need for scalable, high-dimensional imaging-based profiling of cells is rapidly growing as researchers aim to systematically connect molecular signatures to morphological traits, uncover cellular mechanisms, and accelerate drug discovery. High-content morphological profiling has revealed rich phenotypic signatures, especially through fluorescence imaging, which has enabled large-scale screens, pathway reconstruction, and gene-environment studies ^1, 2^. However, fluorescence-based methods are constrained by their reliance on labeling, which limits scalability, introduces phototoxicity, and biases live-cell imaging. Moreover, their throughput is hindered by the trade-off between imaging resolution and field of view (FOV), requiring time-intensive scanning processes to capture large-scale datasets.

Biophysical cytometry offers a label-free alternative by measuring intrinsic properties such as mass, stiffness, and morphology—providing mechanistic insights often missed by molecular assays, and capturing heterogeneity in processes like cell cycle, aging, cancer progression, and drug response ^3^ Quantitative phase imaging (QPI), a cornerstone of biophysical cytometry, enables detailed, label-free analysis of cellular structure and dynamics. Recent studies have demonstrated the utility of QPI in high-throughput drug screening by quantifying parameters such as drug sensitivity, cytotoxicity, and heterogeneity at the single-cell level ^4, 5^. However, current QPI systems are still limited in their ability to extract high-dimensional features at scale, restricting comprehensive profiling of complex cellular phenotypes and their responses to genetic or chemical perturbations.

To overcome these challenges, we introduce a new integrative approach for biophysical cytometry that enables in-depth intrinsic morphological (InMorph) profiling by using a High-Speed Spinning Arrayed Disk (SpAD) imaging platform. SpAD features a continuous, high-speed circular line-scanning mechanism using a spinning disk patterned with radially symmetric cell chambers, unlike conventional microplate readers that rely on slower, raster-scanned rectangular arrays. Operating at speeds over 1000 rpm and coupled with ultrafast QPI, SpAD achieves live-cell imaging at an unprecedented scale and speed—capturing label-free, single-cell morphological data across a large area of ∼100 cm^2^ at sub-cellular resolution. Assays can be completed more than 20 times faster than state-of-the-art fluorescence-based methods, while eliminating the need for labor-intensive labeling ^6^.

SpAD is compatible with standard cell culture workflows and maintains cell health and gene expression, enabling systematic, high-content biophysical profiling across large cell populations. It captures phenotypes from bulk to subcellular levels with exceptional speed and accuracy. We demonstrate that SpAD-powered InMorph profiling enables sensitive, high-throughput screening of both drug and CRISPR-based genetic perturbations, bridging biophysical and molecular phenotypes at a previously unattainable scale and depth. This technology represents a paradigm shift in morphological profiling, unlocking new dimensions of cellular analysis for next-generation, label-free high-content screening.

## Results

### High-throughput On-the-fly SpAD Imaging System

Our platform integrates two key technologies: an ultrafast QPI microscope and a circular Spinning Arrayed Disk (SpAD) optofluidic platform featuring 96 individual imaging chambers. At its core, the system leverages the inherent compatibility between the SpAD’s unidirectional spinning motion and the ultrafast laser line-scanning readout provided by multi-ATOM— a QPI modality that offers at least 100 times faster than the conventional methods without sacrificing resolution ^7–9^ (**Fig. 1a, Supp. Fig. 1**). By synchronizing radial disk translation with high-speed spinning, SpAD achieves real-time, subcellular-resolution imaging across an ultralarge field-of-view (FOV) spanning tens of cm², overcoming the instability and speed limitation of traditional raster scanning.

**Figure 1.**
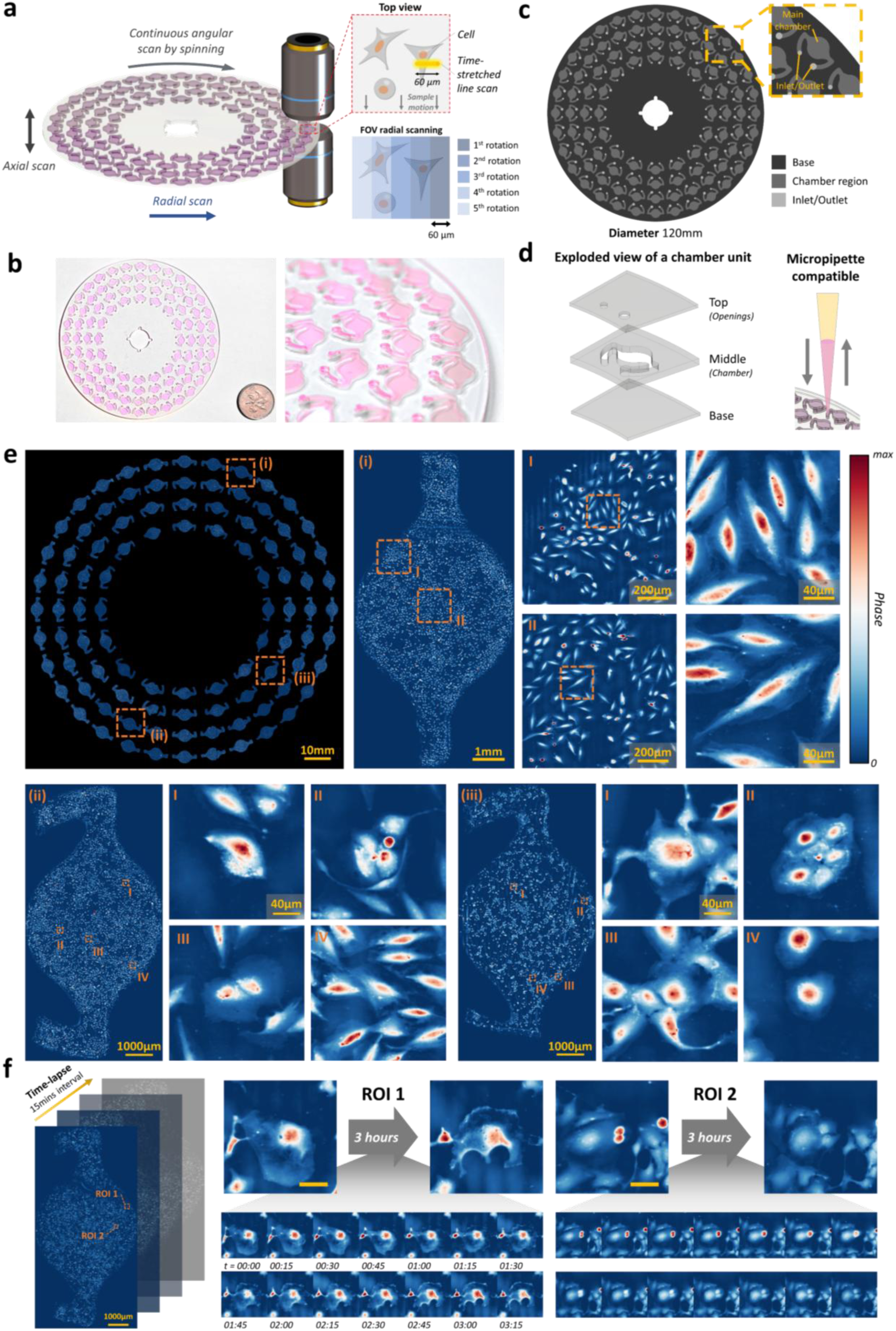
Overview of the SpAD high-throughput imaging system. **(a)** Schematic of the SpAD scanning mechanism. The sample substrate spins continuously, allowing a stationary time-stretched line scan (60 μm in length) to interrogate the sample. Combined radial and axial translation of the spinning substrate enable full coverage of the entire substrate. **(b)** Photograph of the sample substrate containing culture medium. A Hong Kong five-dollar coin (27 mm diameter) is placed adjacent for scale. **(c)** Design layout of the spinning substrate, with chambers arranged in concentric rings. Inset (orange dashed box) illustrates the detailed geometry of an individual chamber. **(d)** Exploded view of a chamber unit, highlighting the three-layered substrate structure that defines the base, chamber walls, and access openings. **(e)** Full-field QPI image acquired from the entire substrate (12 cm diameter) using the SpAD system. Two NSCLC cell lines (H1975 and H2170) were seeded in this example. Non-chamber regions are masked away in black for clarity. Panels (i), (ii), and (iii) show magnified views of individual chambers at different radial positions; chambers (i) and (ii) contain H1975 cells, while chamber (iii) contains H2170 cells. Further enlarged regions (I–IV) demonstrate consistent, high-resolution (∼1 μm) imaging quality across the substrate. **(f)** Time-lapse imaging of live-cell dynamics. In region of interest (ROI) 1, a migrating cell extends its leading edge upward while retracting its trailing edge. In ROI 2, two adjacent cells progressively adhere to the substrate; based on their size and proximity, these are likely daughter cells from a recent mitotic event.

To ensure the robustness of such a high-speed and large-scale imaging capability, further SpAD-specific hardware and software engineering strategies have been integrated. First, we employed continuous axial SpAD scanning to compensate for minor axial image focus drift/jitter during high-speed spinning, and thus to ensure image planes with the best focus can be captured across all 96 chambers (**Methods**). Second, we developed a robust, fully automated image reconstruction pipeline that adaptively scales, aligns, and stitches over 13,000 in-focus image stripes for subsequent chamber-by-chamber QPI reconstructions (**Methods**), resulting in gigapixel to terabyte-scale datasets per SpAD run. Third, we also developed a synchronized real-time visualization tool in the pipeline that allows live image reconstruction and adaptive focus tuning, allowing seamless visual inspection of imaging performance and targeted zooming into regions of interest during image acquisition (**Supp. Fig. 2**).

The SpAD’s design further enables high-throughput parallelization (**Fig. 1b**). In a SpAD, 96 chambers are arranged in a circular asymmetric geometry on a disk to allow parallelized cell assays (**Fig. 1c**). The disk consists of three layers of fused silica wafers: a base layer, a middle chamber layer for the cells seeding and culture, and a top layer with inlet/outlet openings (**Fig. 1d, Supp. Fig. 3**). Any culturing solutions or cell samples can be accessed through the openings with common micropipettes (See next section for compatibility of SpAD with the standard culture protocols).

To demonstrate SpAD’s massive imaging scale and resolution, we seeded non-small cell lung cancer (NSCLC) lines across all chambers and performed QPI imaging (**Fig. 1e**). Across the whole sample disk, several chambers (i, ii and iii) are selected and enlarged in several stages (I-IV) to show the sub-cellular resolution. We visualize chambers at different radii and regions at the chamber edge and centre to show the image quality consistency across the whole disk, critical for downstream quantitative analyses. We emphasize that the phase sensitivity of multi-ATOM in this system is sufficient to reveal the sheet-like lamellipodia as in (ii, III) and (iii, IV). In addition, the whole SpAD imaging FOV (>100 cm^2^) was captured within only minutes - thanks to its high-speed spinning operation. This imaging throughput is particularly advantageous for large-scale time-lapse imaging that uncovers dynamics of rare cellular events, such as cell migration (**Fig. 1f, ROI1**) and mitosis (**Fig. 1f, ROI2**).

### Mechanical stability and cell culture compatibility of SpAD

The SpAD chambers and disk geometry are designed to enable practical, high-throughput live-cell imaging at scale. They feature robust liquid handling, superior mechanical stability, and optimized compatibility with cell culture. To prevent bubble entrapment—a common issue in microfluidic devices— we positioned inlet and outlet channels on opposite sides of the chamber’s circular main region and avoided sharp turns. This design ensures smooth, uninterrupted liquid advancement during pipetting **(Fig. 2a-i)**, resulting in reliable and uniform chamber filling.

**Figure 2.**
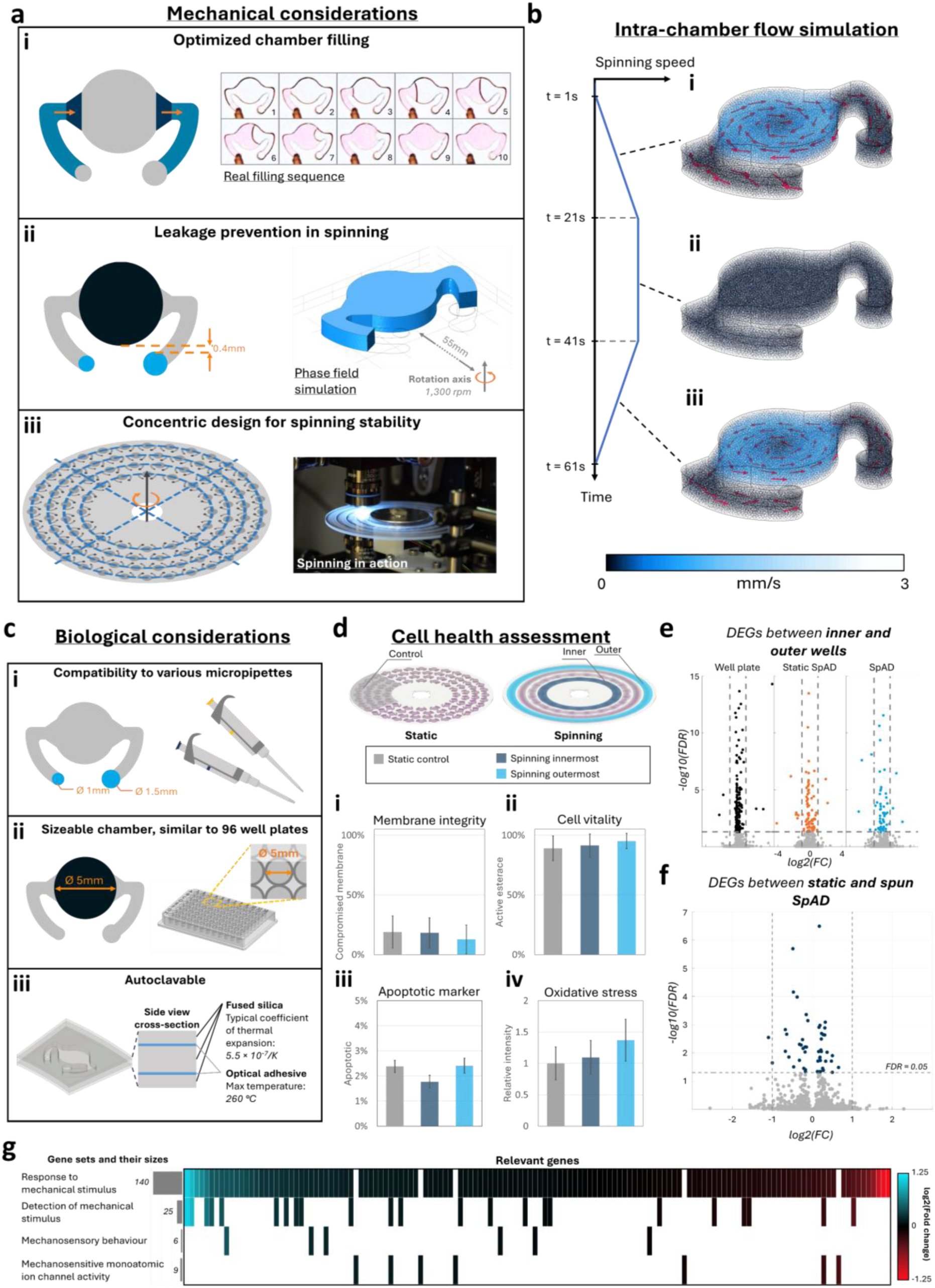
Design rationale and validation of the SpAD sample disk. **(a)** Mechanical design considerations. *(i)* Chamber geometry was optimized to facilitate efficient liquid handling. Opposing inlet and outlet channels, flanked by slanted buffer zones, ensure smooth advancement of the liquid interface during pipetting. Real-life imaging of a chamber-filling sequence demonstrates consistent interface progression. *(ii)* To prevent spillage during high-speed spinning, chamber openings were positioned close to the rotational center. Phase-field simulations confirm that even at 1,300 rpm, fluid remains confined within the chamber. *(iii)* A concentric chamber layout ensures balanced mass distribution, improving rotational stability. **(b)** Intrachamber flow dynamics during operation. Computational fluid dynamics (CFD) simulations modeled fluid motion in a chamber placed at the plate’s periphery under a spin profile comprising acceleration, steady-state spinning, and deceleration (each lasting 20 s). Vortices are transiently observed during acceleration and deceleration; fluid remains largely quiescent during steady rotation, minimizing shear stress on cells. **(c)** Biological design features. *(i)* Asymmetrical opening sizes accommodate both standard and narrow pipette tips, streamlining liquid exchange. *(ii)* The main chamber diameter (5 mm) mirrors that of conventional 96-well plates, facilitating scalability and compatibility. *(iii)* The device supports repeated sterilization by autoclaving and a rapid cleaning protocol, allowing for reliable reuse. **(d)** Assessment of biocompatibility. H2170 spun for 1 hour in SpAD chambers exhibits comparable viability to static controls, validating the platform’s suitability for live-cell assays. **(e)** Volcano plots comparing gene expression between inner and outer wells of a 96-well plate under static SpAD and spun SpAD conditions (left to right). Dotted lines denote significance thresholds: horizontal lines indicate a false discovery rate (FDR) of 0.05, and vertical lines indicate log₂(fold change) cutoffs of ±1. **(f)** Volcano plot comparing static versus spun SpAD conditions. **(g)** Heatmap of genes associated with selected gene ontology (GO) terms related to mechanical stimulus. Colors represent log₂(fold change) values, with genes sorted accordingly. Gray bar plots on the left indicate the number of genes in each GO term.

A central challenge for high-speed SpAD imaging is maintaining media containment during rapid spinning (>1000 rpm), as centrifugal forces could otherwise expel liquid from open chambers. Traditional sealing methods restrict the necessary gas exchange for live-cell assays. To resolve this, SpAD introduces a recessed “pocket” design, placing access openings nearer the rotational axis (**Fig. 2a-ii**). 3D phase-field simulations confirmed the effectiveness of this layout in retaining fluids during operation.

Recognizing that imaging quality is highly susceptible to rotational stability, we engineered the sample disk to spin about its axis of maximal principal inertia by enforcing strict concentric chamber placement and uniform mass distribution (**Fig. 2a-iii**). This design minimizes wobble, preserving imaging focus and resolution (**Supp. Fig. 4**). In addition, the flat chamber roof and floor enable aberration-free, wide-field transmissive imaging, outperforming standard commercial well plates.

Practical usability and workflow integration are at the heart of SpAD’s design. Access openings are fabricated in two diameters to accommodate various pipette tips (**Fig. 2c-i**). Each 5 mm-diameter chamber supports thousands of cells (**Fig. 2c-ii**), matching the capacity of standard 96-well plates and enabling direct adoption into established screening protocols. Chambers are constructed from high-temperature and pressure-resistant materials, allowing for repeated sterilization by autoclaving (**Fig. 2c-iii**).

### Mechanical and Biological Validation of SpAD

To ensure the integrity of the microenvironment experienced by cells within the spinning architecture of the SpAD system, we investigated the system with computational fluid dynamics (CFD) simulations as well as experimental assays. CFD simulations targeted the outermost chamber ring (55 mm from the rotational axis), monitoring possible shear and flow perturbations during a typical spinning protocol: 20 seconds acceleration, 20 seconds at 1,300 rpm, and 20 seconds deceleration (**Fig. 2b**).

During steady-state spinning — the predominant duration of most experimental protocols— CFD simulations revealed negligible fluid movement and no appreciable shear flow within chambers (**Fig. 2b-ii**). In contrast, brief periods of vortex formation occurred only during acceleration and deceleration due to Euler’s force - with the flow velocity peaking at only ∼3 mm/s at the chamber’s mid-plane (**Fig. 2b-i, iii**), well below levels that could mechanically stimulate cells.

In a fully filled spinning chamber, adherent cells experience the outward centrifugal force, the buoyant force from the surrounding medium, and the adhesion force holding the cell to the substrate. As both the cell and medium experience identical centrifugal acceleration, only the small density difference could yield net force, typically on the order of 1-10 pN —far below the threshold needed to detach a cell — so the mechanical stress imposed by spinning is negligible (**Supp. Notes 1**).

To corroborate the simulation results and assess the biological impact of spinning, we performed a collection of cell health assays targeting membrane integrity, esterase activity, apoptosis, and oxidative stress (**Fig. 2d**). Cells from both the innermost and outermost rings were compared against a static control. Another SpAD disk was incubated under identical conditions without rotation (**Methods**).

Microscopy-based analyses indicated that cells subjected to spinning maintained comparable membrane integrity and esterase activity to the static control (**Methods**, **Fig. 2d-i,ii**). Apoptosis rates remained uniformly low across all conditions (average 2.18%), with the lowest fractions in the innermost ring (1.77%) (**Fig. 2d-iii**). Notably, a modest increase in oxidative stress was observed in both inner and outer regions relative to the control (**Fig. 2d-iv**). All analyses were performed on pooled data from replicate wells to ensure statistical robustness.

To evaluate the consistency of cellular behavior across different chamber locations, we analyzed gene expression differences between cells in the inner and outer wells of the 96-well plate, both under static and spinning SpAD conditions (**Methods**, **Fig. 2e**). Among the 60,725 mapped genes, only 171, 84, and 59 genes were differentially expressed across inner and outer wells in the 96-well plate, static SpAD, and spinning SpAD comparisons, respectively. This minimal number of differentially expressed genes (DEGs) indicates negligible positional impact on gene expression. Furthermore, comparing static versus spinning conditions across all wells revealed only 50 DEGs (**Fig. 2f**), indicating that spinning has negligible transcriptomic impact on the cells. Gene ontology (GO) analysis further revealed no significant changes in mechanical stimulus-related terms, underscoring SpAD is gentle on cells and preserves their intrinsic gene expression signatures (**Fig. 2g**).

Together, these findings establish that SpAD maintains a stable and uniform microenvironment for cells during extended centrifugation, supporting its suitability for sensitive biological assays. Crucially, the design minimizes both well-to-well batch effects ^10^ and spinning-induced mechanical perturbation enabling high-throughput experimentation without compromising cell health or molecular signatures.

### SpAD Single-cell Image-based Biophysical Profiling of Drug Responses

Majority of image-based phenotypic screening approaches rely on fluorescence microscopy ^2^, requiring laborious preparation steps, high reagent costs, and lengthy protocols. Here we demonstrate a streamlined, label-free drug screening pipeline with SpAD that overcomes these limitations. Leveraging high-speed SpAD imaging and in-depth InMorph profiling, the assay time can be shortened by ∼20 times compared to the leading fluorescence imaging method (**Supp. Fig. 5**) without compromising high-content screening.

The SpAD image-based drug screening assay (**Fig. 3a**) utilized two NSCLC cell lines—H1975 (adenocarcinoma, EGFR T790M mutant) and H2170 (squamous cell carcinoma, EGFR wildtype)—chosen for their distinct morphologies and EGFR profiles. Four anticancer drugs with diverse mechanisms were tested: cisplatin (DNA damage), docetaxel (microtubule stabilization), erlotinib (EGFR inhibition), and gemcitabine (DNA chain termination) (**Methods**, **Fig. 3b**). Each drug was screened at five concentrations with replicates, which utilized 88 chambers including controls (**Fig. 3c**; concentrations in **Supp. Fig. 6**; images in **Supp. Fig. 9**). Assay reproducibility was validated using intra-experiment replicates and independent runs on separate days. This demonstrates SpAD’s robust capacity for high-throughput, biophysical drug screening across multiple cell types and drug classes (**Fig. 3d**).

**Figure 3.**
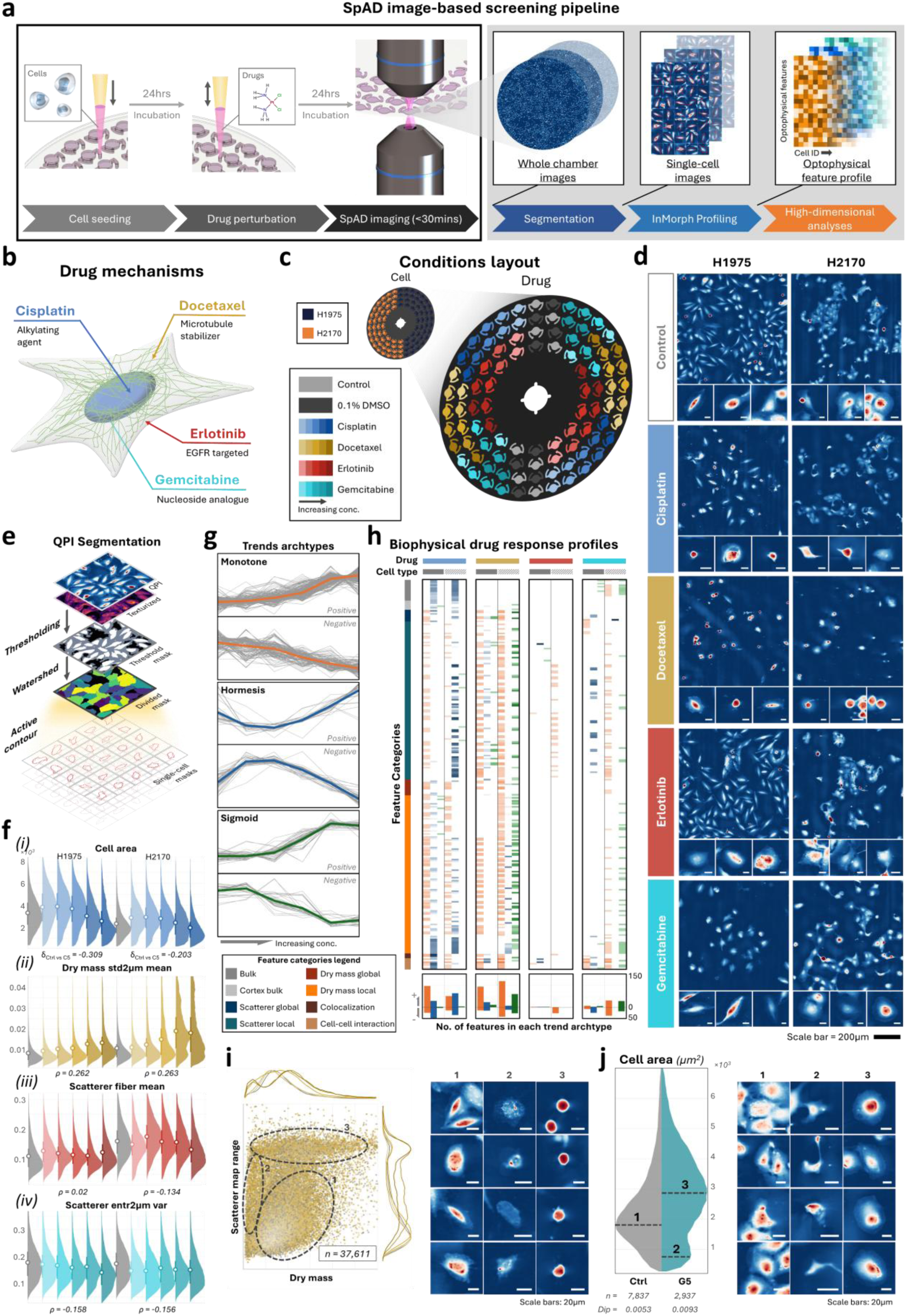
Label-free, image-based drug screening using SpAD. **(a)** Experimental workflow. Cells are seeded and allowed to adhere for 24 h before drug treatment. Following another 24 h incubation, the spinning plate is directly imaged by the SpAD system. Acquired images are analyzed through segmentation, feature extraction, and high-dimensional profiling at the single-cell level. **(b)** Mechanisms of action (MoA) for selected drugs. Cisplatin and gemcitabine exert cytotoxicity via DNA damage, albeit through distinct pathways. Docetaxel disrupts mitosis by hyperstabilizing microtubules, whereas erlotinib inhibits the epidermal growth factor receptor (EGFR). **(c)** Plate layout indicating drug identity and concentration. Each color represents a different compound, with intensity denoting dose (darker shades indicate higher concentrations). **(d)** Representative QPI images of drug-treated and control cells. Left and right columns correspond to H1975 and H2170 cell lines, respectively. Drug-treated samples reflect the highest concentration condition; controls correspond to 0.1% DMSO. The scale bar (black) is located at the bottom. Segmented cells are shown below each image with the corresponding scale bars (white, 20 µm) located at the corner of the image. **(e)** Segmentation pipeline for quantitative phase imaging (QPI). The training-free workflow includes: (i) thresholding of raw and texture-enhanced QPI images, (ii) marker-based watershed segmentation, and (iii) active contour-based refinement to enhance boundary accuracy. **(f)** Dose-dependent feature responses. Histograms display selected morphological and biophysical features across increasing drug concentrations for both cell lines. Mean (white circle) and interquartile range (bars) are shown. Effect sizes and monotonicity are quantified using Cliff’s delta (δ_Ctrl vs C5_) and Spearman’s ρ, respectively. **(g)** Canonical shapes of trend categories. Trends in extracted features are classified into three archetypes: monotonic, hormetic, and sigmoidal. Mean-normalized trends (gray lines) from all features are overlaid for each category, with bold colored curves denoting the average profile. Detailed methodology for trend classification is provided in **Methods**. **(h)** Comprehensive drug response profiles. Columns correspond to drug–cell line pairs; rows represent individual features, color-coded by feature category. Bar plots below each matrix indicate the count and directionality of features within each trend archtype. **(i)** Subpopulation analysis for docetaxel treatment. A two-feature scatter plot reveals three distinct cell subclusters in H1975 cells. Marginal histograms illustrate feature distributions. Example images of each subcluster are shown below. Scale bars: 20 µm. **(j)** Subpopulation emergence under gemcitabine exposure. Bimodal distributions in “*Cell Area*” at high drug concentrations indicate distinct subclusters. Representative images of each peak are displayed. Subpopulation separation is statistically validated using Hartigan’s Dip Test (Dip). Total cell count (n) is listed alongside.

To enable robust single-cell InMorph profiling across diverse perturbations in SpAD, we developed a novel, training-free segmentation algorithm that integrates multiple tailored image-processing strategies (**Fig. 3e**). Unlike the deep learning-based methods that require large, annotated datasets and computational training, our approach exploits the intrinsic morphological cues captured by the pseudo-3D contrast in QPI (**Supp. Fig. 8, Methods**), ensuring strong generalizability across conditions. Building upon feature extraction tools such as CellProfiler ^11^ and our earlier work ^9, 12^, we extracted a set of 377 biophysical features from each segmented cell. These features span bulk morphological descriptors, intracellular textural metrics, spatial distributions of scatterers and dry mass, and intercellular interaction indices (see **Supp. Table** for the full feature list).

This high-dimensional representation enables quantification of subtle cellular responses to pharmacological perturbations. For example, with increasing concentrations of docetaxel, we observed an elevated abundance of 2 µm-sized dry mass structures within cells, as reflected by an upward shift in the QPI feature “*dry mass std2µm mean”* (**Fig. 3f-ii**). In contrast, gemcitabine treatment led to a downward shift in a bright-field feature “*scatterer entr2um var”*, suggesting a more homogeneous spatial distribution of 2-µm-sized scatterers (**Fig. 3f-iv**). Notably, cisplatin elicited a hormetic response in “*cell area”* across both cell lines (**Fig. 3f-i**), potentially reflecting enhanced growth or adhesion at lower concentrations. Differences in drug sensitivity were also apparent between cell types, with the erlotinib-resistant H1975 cells exhibiting minimal response (**Fig. 3f-iii**).

To analyze complex patterns in these biophysical responses, we developed a trend detection algorithm to classify feature behavior across concentration gradients (see **Methods**). We identified three characteristic trend types commonly observed in drug screens ^13^ (**Fig. 3g**). By aggregating trend classifications across drugs, cell types, and features, we constructed a comprehensive biophysical response profile for each drug-cell combination (**Fig. 3h**). After filtering non-responsive features, 258 remained, forming the basis for comparative profiling. These profiles revealed that cisplatin and docetaxel induced the strongest responses, each with distinct feature signatures, consistent with 72-hour MTT assay results (**Supp. Fig. 7**), yet. Cisplatin induced prominent hormetic trends (marked in blue), especially in bulk features, while docetaxel induced more sigmoidal changes (green), particularly in H2170 cells. The nearly blank profile of H1975 in response to erlotinib confirmed its resistant phenotype. Gemcitabine generated moderate responses, consistent with its known time-dependent mechanism of action ^14^. Overall, these profiles highlight the ability of biophysical features to reflect diverse mechanisms of action and cellular susceptibilities.

### Revealing Heterogeneous Single-Cell Biophysical Responses to Drugs

Beyond trend analysis, SpAD’s single-cell resolution enabled us to uncover drug-induced subpopulation dynamics. In H1975 cells treated with docetaxel, three subpopulations emerged based on dry mass and scattering features (**Fig. 3i**): healthy cells which are dominant at low concentrations (subpopulation 1), apoptotic bodies or membrane remnants with very low dry mass (<100 pg) prevalent at high concentrations (subpopulation 2), and rounded, intermediate-mass cells indicative of mitotic or stressed states at high doses (subpopulation 3) (**Fig. 3i**).

For H2170 cells treated with a high dose of gemcitabine (G5), we observed a pronounced bimodal distribution in cell area (**Fig. 3j**). This deviation from the typical unimodality was confirmed using Hartigan’s Dip test ^15^, which was doubled in the test statistic compared to the control. Morphological analysis of cells corresponding to each distribution peak revealed three further distinct subpopulations. Peak 1 included cells morphologically similar to untreated controls, characterized by prominent cytoplasmic cores. Peak 2 consisted of significantly shrunken cells, likely apoptotic in nature ^16^. Intriguingly, Peak 3 represented cells with larger projected areas and prominent lamellipodia. This group emerges only at the highest drug concentration and coincides with reduced cell counts, compared to control (**Supp. Fig. 11**), likely represents a resistant phenotype exhibiting epithelial-to-mesenchymal transition (EMT)-like adaptations, consistent with previous reports linking gemcitabine resistance to increased motility ^17^, and cytoskeletal remodeling ^18^.

### Label-Free CRISPR Screening of Cell Fusion Using Biophysical Features

CRISPR-based functional genomics screens have transformed systematic gene perturbation studies, and when combined with morphological profiling, can reveal how genetic changes affect cell structure and function ^1, 19–22^. However, the impact of genetic perturbations on biophysical morphology (i.e., InMorph) remains largely unexplored. Gaining this knowledge would complement current fluorescence-based profiling and provide new insights into the roles of biophysical traits in cell states and functions. Here, we explore a novel application of our SpAD imaging platform to perform fully label-free live-cell CRISPR screening, using syncytia formation—induced by the SARS-CoV-2 spike protein—as the phenotypic endpoint.

To model this, HEK293T cells transfected with SARS-CoV-2 spike (referred to as “sender cells”) were co-cultured with ACE2-expressing recipient cells harboring gene knockouts. Within 24 hours, susceptible recipient cells fused into multinucleated syncytia, a hallmark of spike-mediated viral pathogenesis (**Methods**) ^23^. Unlike the previous split-GFP systems that require genetic reporters and fixation protocols ^24, 25^, SpAD’s QPI and brightfield imaging directly visualize fusion events in live cells, accelerating assay speed and eliminating labeling steps.

To test our approach, we targeted four gene knockouts—ACE2, AP2M1, FCHO2, and CAB39—all previously shown to reduce syncytia formation. Among them, ACE2 knockout, the direct receptor of spike, showed the most pronounced effect ^26^. Following SpAD imaging, a subset of chambers was also imaged using fluorescence microscopy to provide ground-truth syncytia labels (**Fig. 4a**). These dual-modality datasets were co-registered (**Fig. 4b**; **Methods**) to build an annotated training set based on both QPI and fluorescence. Considering the morphological complexity of syncytia, analysis was focused on cell clusters, from which we identified two main subpopulations based on the InMorph profile (**Fig. 4c**): flattened and rounded clusters, as confirmed by image inspection (**Supp. Fig. 12b**). Most syncytia (79.0%) were found in the flattened group, which was selected based on the distinctive dry mass distribution features for downstream analysis (**Supp. Fig. 12a**).

**Figure 4.**
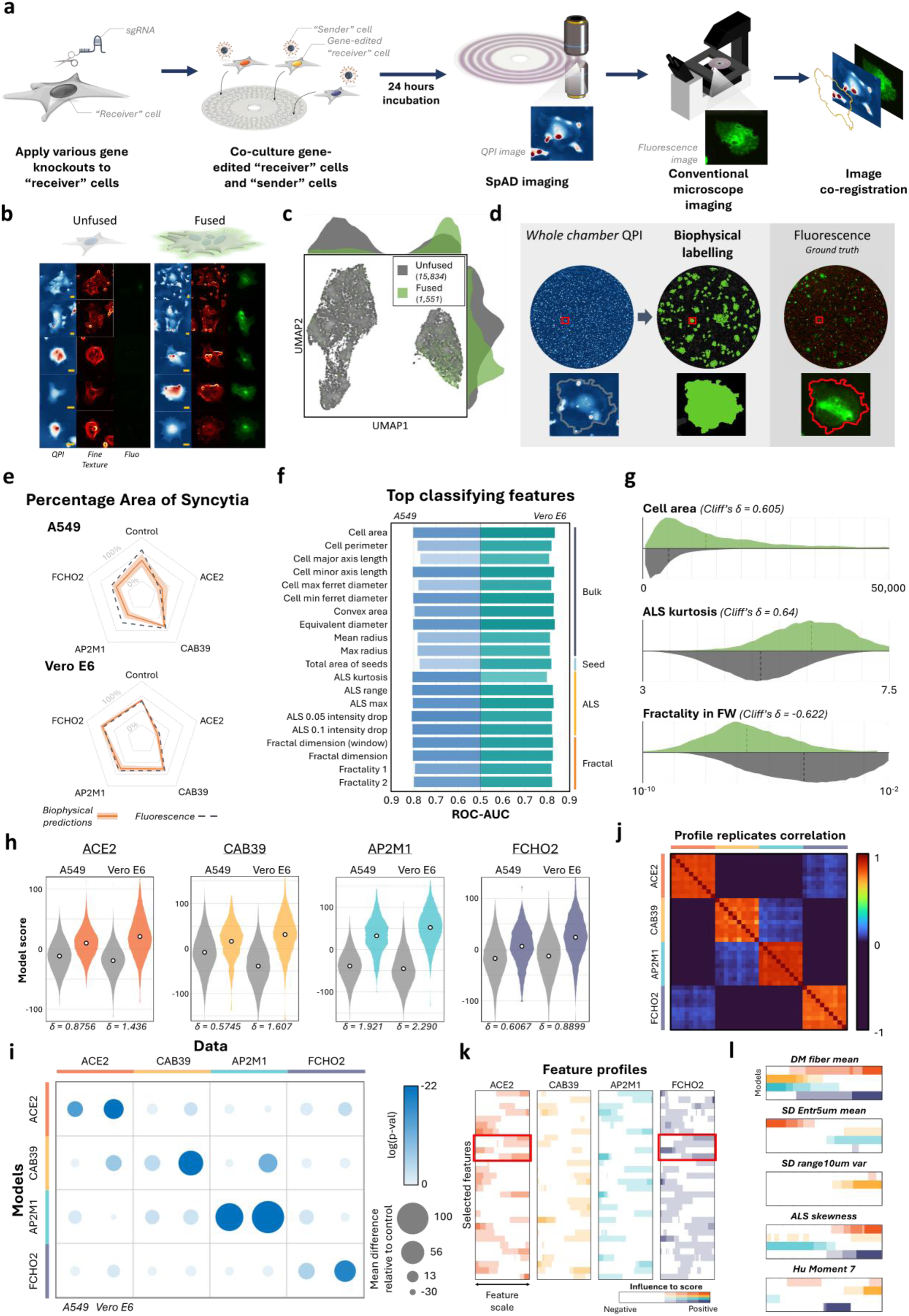
Label-free, image-based CRISPR screening for SARS-CoV-2 spike-induced cell fusion. **(a)** Overview of the experimental workflow. Gene knockout “receiver” cells are co-cultured with SARS-CoV-2 spike-expressing “sender” cells for 24 h to induce fusion. Prior to conventional fluorescence imaging, quantitative phase imaging (QPI) is performed using the SpAD system. Fluorescence and QPI data are subsequently co-registered for downstream analysis. **(b)** Representative images of unfused cells and fused syncytia. Co-registered fluorescence images (green signal from split-GFP system indicates fusion), QPI, and texture-enhanced contrast images are shown side-by-side. Scale bar, 20μm. **(c)** UMAP projection of A549 colonies using extracted biophysical features. Color-coded clusters correspond to distinct morphological populations (legend); numbers in parentheses indicate colony counts. The left cluster comprises spherical cells, while the right contains adherent colonies. Projected histograms indicate that syncytia predominantly localize to the adherent colony cluster (see **Supp. Fig. 12** for more examples). **(d)** Syncytia labeling via QPI and a machine learning classifier trained on co-registered fluorescence labels. The model enables biophysical annotation of syncytia in the absence of fluorescence. **(e)** Comparison of syncytia occupied areas across gene knockouts and cell lines using biophysical predictions and fluorescence-based measurements. Orange lines and shaded regions indicate the mean and standard deviation of the biophysical predictions, respectively. **(f)** Top features discriminating between unfused colonies and syncytia, ranked by area under the receiver operating characteristic curve (ROC-AUC). Left bars: A549-ACE2 cells; right bars: Vero E6-TMPRSS2 cells. Feature category annotations are shown on the right. **(g)** Distributions of selected discriminative features. Means are denoted by dashed lines; Cliff’s delta (δ) values quantify effect sizes between syncytia and unfused colonies. Note: x-axis of “Fractility in FW” is log-scaled. **(h)** Gene-specific colony classification using generalized additive models (GAMs). For each gene knockout, a separate GAM is trained to distinguish knockout from control colonies in both A549 and Vero E6 cell lines. Violin plots show score distributions; Cliff’s δ quantifies separation. **(i)** Specificity of gene-knockout classification. GAMs trained on individual knockouts are evaluated against all other knockouts. Circle size and color represent mean score difference and log-transformed p-value (Wilcoxon rank-sum test), respectively. **(j)** Model stability across replicates. Pearson correlation matrix of model outputs from 10-fold cross-validation shows consistency among trained models. **(k)** Condensed partial dependence (PD) profiles for selected features across models. X-axis: normalized feature values; Y-axis: selected features. Color intensity indicates feature influence on model output; red boxes highlight conserved feature effects across multiple models. **(l)** Direct comparison of partial dependence profiles. Models are arranged side-by-side to visualize differential feature responses across knockouts.

Using the annotated dataset, we trained a supervised machine learning model to classify syncytia versus unfused clusters (**Methods**). The model was subsequently applied across the entire dataset—including the chambers without the fluorescence data—in order to test the syncytia predictions based solely on label-free imaging (**Fig. 4d**). Across cell lines and knockout conditions, syncytia area estimates derived from SpAD-based predictions closely mirrored those obtained from fluorescence imaging (**Fig. 4e**). This demonstrates that biophysical imaging alone is sufficient to capture fusion phenotypes with comparable accuracy, while markedly simplifying experimental workflows.

Feature analysis showed that 19 of the top 20 discriminative InMorph features were consistent across cell lines, dominated by size-related metrics (**Fig. 4f, Supp. Fig. 13**), with many exceeding AUC values of 0.8. Beyond cell size, single cell textural features derived from angular light scattering (ALS) and single-cell fractal geometry have been proven informative for morphological profiling ^12^. For instance, more significant large-angle light scattering from single cells, which is attributable to increased subcellular granularity (as quantified by the feature ALS kurtosis) ^12^, is observed in the fused colonies (effect size = 0.64) (**Fig. 4g-ii**). In contrast, fused syncytia exhibited reduced deviation from fractality (effect size = 0.622; **Fig. 4g-iii**), indicating a more fractal internal structure compared to unfused cell clusters. Collectively, these findings not only enable accurate syncytia classification, but also provide mechanistic insights into structural transformations during fusion.

### Gene Knockouts Induce Distinct Label-Free Phenotypes

We next investigated whether gene-specific knockout effects could be discerned through label-free SpAD imaging and InMorph profiling. Focusing on unfused cell populations— identified using the pre-trained syncytia classifier—we trained a series of generalized additive models (GAMs) to predict the likelihood of individual gene knockouts based on biophysical features (**Method**). Both A549 and Vero E6 cell lines were included in the training to promote model generalizability.

Each model effectively discriminated its target knockout from control populations in cross-validation (**Fig. 4h**). The AP2M1 knockout, in particular, exhibited a striking separation (mean effect size = 2.11), while even the weakest-performing model, targeting CAB39 in A549 cells (effect size of 0.5745). Specificity tests confirmed that each model scored highest on its intended knockout population compared to others (**Fig. 4i**), and 10-fold cross-validation revealed strong reproducibility across model replicates (**Fig. 4j**).

Feature-level visualization using partial dependence plots (PDPs) revealed distinct biophysical signatures for each gene knockout **(Fig. 4k**), with consistent patterns across 10 replicates supporting the robustness of the learned models (**Supp. Fig. 14**). Interestingly, certain features exhibited overlapping dependencies across multiple knockouts (highlighted box, **Fig. 4k**), suggesting that shared phenotypic programs may be triggered by different gene perturbations.

To further dissect the basis of model discrimination, we examined the PDPs of individual biophysical features across different gene knockout models (**Fig. 4l**). These PDPs revealed that certain features contribute to distinct subsets of knockouts in divergent ways. For instance, *DM Fiber Mean* was positively associated with ACE2 and FCHO2 knockouts, while its lower values were more predictive of CAB39 and AP2M1 perturbations. Other features exhibited more selective associations—*SD Range10um Var* indicated that CAB39 knockouts had greater variability in local contrasts, whereas *Hu Moment 7* revealed lower cell shape symmetry in CAB39 knockouts and higher cell shape symmetry in FCHO2 knockouts. Importantly, these features are informative only in combination in order to classify knockouts, reflecting the nonlinear and multifactorial nature of the models. Together, these results demonstrate that interpretable machine learning can map gene-specific biophysical phenotypes from label-free imaging data alone.

## Discussion

Imaging-based phenotyping is central to high-content screening but remains limited by complex labeling protocols and slow imaging throughput. These constraints hinder large-scale studies and add significant labor, cost, and batch variability. To overcome these barriers, we introduce SpAD—a high-throughput, label-free imaging platform that combines continuous circular scanning, ultrafast QPI, and a novel circular sample array.

On the hardware front, unidirectional spinning—replaces traditional raster scanning, eliminating motion artifacts and mechanical instability. Its concentric 96-well arrangement supports robust, culture-compatible sample handling. Integrated with multi-ATOM, an ultrafast QPI modality, SpAD simultaneously acquires brightfield and phase information without the need for labeling.

Altogether, this platform achieves subcellular-resolution imaging across >200,000 single cells per run, with the entire assay workflow completed in under 15 minutes— a ∼20-fold reduction in experimental time compared to fluorescence imaging-based methods.

On the analytical front, InMorph Profiling extracts rich, multidimensional biophysical features from each cell, enabling deep phenotypic characterization. Further powered by machine learning-based analysis, this high-throughput, high-resolution approach enables rapid, label-free phenotyping at scale. In drug screens covering 330,000 cells, InMorph profiling using SpAD sensitively detected drug-induced phenotypic changes, resistant subpopulations all without labeling. More significantly, we demonstrated that SpAD imaging assay is able to classify the gene knockouts that impact viral syncytia, solely based on distinct InMorph profiles. The platform’s ability to uncover distinct biophysical signatures from brightfield and phase contrast - all without the need of any labelling - opens new possibilities for large-scale, label-free subpopulation analysis.

Unlike imaging-based screening methods that rely on fluorescence labeling—such as Cell Painting ^27^ and VIBRANT ^28^—SpAD eliminates time-consuming and labor-intensive sample preparation steps. Fluorescent labeling not only increases assay complexity but also introduces sources of batch effect ^29^. Its label-free, ultrafast imaging streamlines workflows and scales seamlessly, all while maintaining compatibility with standard cell culture and handling protocols. While SpAD does not inherently provide molecular specificity, our results show that even straightforward machine learning models can extract biologically meaningful insights from its high-content, label-free data (**Fig. 3**–**4**).

SpAD’s scalable and automated approach is ideally suited for generating large datasets for downstream AI-driven analysis. Its non-invasive nature allows direct integration with other single-cell techniques, such as RNA sequencing ^30^ and mass spectrometry ^31^, to build comprehensive multi-omic atlases by correlating morphology with molecular profiles. In translational contexts, SpAD’s speed and low labor demands enable rapid, scalable precision medicine screens—particularly valuable for patient-derived cells and clinical applications.

Despite these advances, some limitations remain. SpAD’s parallelization is currently limited by manual liquid handling; scaling to higher-density formats (e.g., 384-well plates) would benefit from automation. While axial scanning improves focus, it lengthens imaging time; future improvements in spinning precision or computational refocusing ^32^ could alleviate this trade-off. SpAD currently prioritizes throughput over longitudinal observation, but integrating a custom incubator may enable dynamic live-cell studies. Looking ahead, future work will expand the repertoire of gene knockouts, drug perturbations, and cell types, moving toward a comprehensive atlas of label-free morphological signatures. Its high dimensionality and throughput make SpAD a powerful data engine for deep learning ^33^, positioning it to bridge image-based phenotyping with actionable functional insights and accelerate next-generation precision medicine ^34^.

## Acknowledgement

We thank the Centre of PanorOmics Sciences (CPOS), University of Hong Kong for their expert opinions and sequencing services for our transcriptomic profile experiment. The work is supported by Advanced Biomedical Instrumentation Center, the Research Grants Council (grant no. 17125121, 14125924, RFS2021-7S06), the Innovation and Technology Commission of the Hong Kong Special Administrative Region of China (grant no. ITS/318/22FP, ITS/408/23FP), Platform Technology Funding of the University of Hong Kong, and the Centre for Oncology and Immunology Limited under the Health@InnoHK Initiative funded by the Innovation and Technology Commission, The Government of Hong Kong SAR, China.

## Online Methods

### SpAD imaging system

The SpAD imaging platform is built upon the ultrafast, laser-scanning modality known as Multi-ATOM ^7, 9^, optimized for high-throughput, label-free phase imaging. A time-stretched line-scan is generated by coupling a 10 MHz mode-locked laser into a single-mode fiber spool with a group velocity dispersion of 1.78 ns/nm, followed by spectral dispersion through a diffraction grating (Thorlabs, GP3508P). This produces a temporally encoded 1D line-scan, which is relayed and focused onto the sample via a lens assembly and a matched pair of objective lenses (Olympus LCPlan N 50×), yielding an illumination line of approximately 60 µm in length and a numerical aperture of ∼0.56.

After interacting with the sample, the modulated beam is recombined using the same diffraction grating and split into four replicas. Each replica is partially occluded from a different direction to encode phase-gradient information, forming four directionally distinct differential phase contrast (DPC) images. These replicas are time-multiplexed through optical paths of varying fiber lengths and detected by a high-bandwidth photodetector (Newport, 1544-B). From the phase-gradient signals, both brightfield and quantitative phase images are reconstructed in post-processing. A complete system schematic is provided in **Supp. Fig. 1**.

### Real-time visualization and focus tuning

To enable optimal imaging performance, the SpAD system integrates real-time visualization and interactive focus tuning through a trigger-based image reconstruction module embedded within the FPGA-based data acquisition framework. Image acquisition is synchronized to the rotation of the sample disk via a Hall sensor signal, which initiates each acquisition cycle. The acquisition duration is dynamically determined based on the measured spinning period to ensure complete coverage of a full revolution.

During each cycle, a full revolution of 1D line-scan data is captured. Clock-level synchronization between the FPGA and the pulsed seed laser allows these sequential 1D scans to be rapidly assembled into 2D image frames. These frames are rendered in real time at a refresh rate corresponding to the disk’s rotation frequency, enabling intuitive sample navigation and live focus adjustment.

For detailed inspection of specific areas, users can define regions of interest using polar coordinates. The system then applies a programmable delay relative to the acquisition trigger and adjusts the translation stage position accordingly, enabling targeted, real-time zooming into selected regions.

### Robust, Automated, and High-Throughput Correlative Stitching

To reconstruct high-resolution composite images of all 96 chambers (each 5 mm in diameter) from narrow optical sections (∼60 µm in width), a robust and fully automated image reconstruction pipeline was developed. The SpAD system captured complete circular rings of DPC images, which were segmented into short image stripes based on the known spatial layout of the chambers.

To compensate for radial scanning speed variations inherent to the rotating acquisition geometry, adaptive scaling was applied to each stripe. Axial focus drift was addressed via Z-stack scanning, enabling selection of the optimal focal plane for each chamber individually. Within each chamber, adjacent image stripes were stitched together using cross-correlation of overlapping regions, ensuring consistent alignment and seamless phase reconstruction across the ultrawide field. This stitching algorithm also accounted for minor fluctuations in rotation speed and mechanical jitter. In a typical experiment, the pipeline executed over 13,000 stitching operations error-free, demonstrating its robustness and scalability for high-throughput imaging applications.

### SpAD sample disk fabrication

Patterned fused silica wafers of varying thicknesses (AlfaQuartz) were sequentially cleaned with deionized (DI) water, acetone, and isopropanol (IPA) to remove surface contaminants. To bond the middle and bottom quantz layers, a thin film of UV-curable optical adhesive (Norland Optical Adhesive, NOA 61) was applied to the middle layer using a manual kiss-coating technique. Briefly, a small amount of adhesive was deposited onto a glass slab and evenly spread using a soft brush. The perforated middle layer quantz was then placed onto the adhesive-coated slab and gently dragged across to achieve a uniform, thin coating. This layer was subsequently aligned and attached to the bottom wafer. If air gaps were present at the interface, additional adhesive was applied to the perimeter, allowing capillary action to fill the voids. To ensure uniform adhesion and minimize warping, the assembly was sandwiched between two 1 cm-thick square glass slabs and clamped at all four corners using binder clips to apply even pressure. A pre-curing step was performed using a 60W-rated 365nm UV torch for 1 minute while the assembly remained between the slabs. To remove excess adhesive, the assembly was immersed in acetone for 10 s with gentle agitation, followed by immediate immersion in DI water to halt adhesive dissolution and prevent overexposure. After a final rinse in IPA and drying with lint-free tissue, the top layer was attached using the same procedure. A full UV curing was then performed by illuminating the entire assembly for at least 10 min. The assembled disk was aged at room temperature for one week prior to experimental use to allow complete stabilization of the adhesive.

### SpAD sample disk experiment preparation

Previously used disks were first decontaminated by immersion in 25% bleach solution. The chambers were then filled with RIPA lysis buffer (ThermoFisher Scientific) and incubated for 15 minutes to dissolve any residual cellular or proteinaceous material. Between each solution exchange, the chambers were thoroughly rinsed with deionized (DI) water to remove residual reagents. Sequential immersions in acetone and isopropanol (IPA) were subsequently performed to eliminate remaining contaminants.

Before initiating new experiments, the cleaned disk was sterilized by autoclaving and dried overnight in a 70 °C oven. To promote cell adhesion, the chamber surfaces were coated with fibronectin. The chambers were first wetted with DI water to ensure uniform surface hydration. A fibronectin working solution (40 µg/mL) was prepared by diluting stock fibronectin (Sigma-Aldrich) in DI water. The wetted chambers were then filled with the fibronectin solution and incubated at 37 °C for at least 1 hour. Following incubation, the solution was aspirated, and the chambers were rinsed once with the intended cell culture medium to remove excess unbound protein. The medium was then removed, leaving the disk ready for immediate cell seeding.

### Computational Fluid Dynamics (CFD) Simulation

All simulations were conducted using COMSOL Multiphysics 5.5. For the intrachamber flow analysis, the *Laminar Flow* physics interface was used, assuming steady-state, incompressible flow. Water was modelled using COMSOL’s default material properties. The chamber was located 55 mm from the centre of rotation, and all chamber walls were defined with a no-slip boundary condition. A pressure boundary condition (0 Pa) was applied at the chamber openings to approximate exchange with the surrounding environment.

To simulate spinning conditions, the rotating frame of reference was implemented by defining centrifugal, Coriolis, and Euler forces as volume forces in the momentum equations. Angular velocity was time-dependent and defined according to the profile shown in **Fig. 2b**. The governing equations were solved under time-dependent conditions. A physics-controlled mesh with *fine* element size was used, with refinement near boundaries and corners to ensure accuracy in regions of high shear or velocity gradients.

For multiphase simulations, the *Two-Phase Flow, Phase Field* interface was used to model air–liquid interactions near the chamber openings. Initial conditions assigned air regions at the openings and liquid (water) in the rest of the chamber. Open boundary conditions were applied to both inlet and outlet to permit interface deformation and fluid exchange. The same rotating frame and chamber geometry were used as in the laminar flow model. The interface thickness parameter and mobility were set to 2.63 × 10⁻³ and 1.0 × 10⁻⁵, respectively, to resolve interface dynamics.

### Assessment of Cell Health on the SpAD Platform

Two types of SpAD sample disks— Disk_spin_ and Disk_static_—were prepared using the procedure outlined above. H2170, a human non-small cell lung cancer (NSCLC) cell line (ATCC), was cultured in RPMI-1640 medium with ATCC modifications (ThermoFisher Scientific) under standard incubator conditions (37 °C, 5% CO₂). Subculturing was performed two to three times per week using 0.25% Trypsin-EDTA (ThermoFisher Scientific). For seeding, trypsinized cells were resuspended in fresh medium and adjusted to a final concentration of 1.5 × 10^5^ cells/mL. A 40 µL aliquot of the suspension was dispensed into the outermost ring and innermost chambers of Disk_spin_, and into a representative segment of chambers in Disk_static_. A total of 12 chambers per condition were seeded to ensure sufficient statistical power. Disks were incubated overnight at 37 °C to facilitate cell attachment, with a humidifying water reservoir placed underneath to prevent media evaporation. Following attachment, Disk_static_ was left at room temperature for 1 hour, while Disk_spin_ was mounted onto the SpAD system and subjected to 1 hour of rotation at 1,300 rpm.

To assess cell health, four chambers per condition were stained with calcein-AM and ethidium homodimer-1 (ThermoFisher Scientific, L3224) for viability and membrane integrity, four with a caspase 3/7 detection reagent (ThermoFisher Scientific, C10723) for apoptosis, and four with CellROX Orange Reagent (ThermoFisher Scientific, C10443) for oxidative stress, all following manufacturer protocols. Reagent volumes were scaled appropriately for the 40 µL working volume of the SpAD chambers, as opposed to standard 96-well plate volumes. Disks were subsequently imaged using a commercial widefield fluorescence microscope equipped with matched filters and brightfield optics. Images were analyzed using custom MATLAB scripts. Brightfield images were segmented using a texture-based algorithm to estimate total cell count per grid (355 µm × 355 µm), while fluorescence images were thresholded to detect signal-positive cells. Health metrics were computed as the percentage of apoptotic, metabolically active, or membrane-compromised cells, depending on the stain used. For oxidative stress analysis, the amount of reactive oxygen species was reported as a ratio of cellular fluorescence signal to background.

### Differential gene expressions (DEG) analysis

To evaluate platform-wide consistency and potential transcriptional effects of spinning, we performed next-generation sequencing (NGS) to profile transcriptomes of H1975 cells under different experimental conditions. Cells were seeded into six groups: inner and outer wells of a standard 96-well plate, inner and outer wells of static SpAD, and inner and outer wells of spinning SpAD. After 24 hours of attachment and growth, the 96-well plate and static SpAD groups were left at room temperature for 1 hour, while the spinning SpAD group was subjected to 1,300 rpm rotation for 1 hour. Cells were then harvested using trypsin, and total RNA was extracted using the RNeasy Mini Kit (QIAGEN, 74104) according to the manufacturer’s protocol. Each condition was performed in triplicate.

Sequencing was conducted on an Illumina NovaSeq 6000 platform. Raw reads were filtered to remove low-quality sequences and reads shorter than 40 bp. Filtered reads were aligned to the human reference genome (GRCh38) using STAR v2.7.8 with default parameters. Transcript abundance was quantified using RSEM v1.2.31, and differential gene expression analysis was performed with EBSeq v2.0.0. Genes with a false discovery rate (FDR) below 0.05 were considered significantly differentially expressed.

### SpAD drug screening

SpAD sample disks were prepared and seeded following the standardized protocol described previously. The test involved two NSCLC cell lines—H1975 (adenocarcinoma) and H2170 (squamous cell carcinoma)—which exhibit distinct morphologies and EGFR expression profiles (wildtype in H2170 and T790M mutation in H1975). Both cell lines were cultured and maintained under the same conditions. Cells were seeded and incubated for 24 hours, followed by drug treatment for another 24 hours before SpAD imaging (**Fig. 3a**).

Four anticancer drugs, with diverse mechanisms of action were selected to demonstrate the ability of SpAD to capture distinct drug-induced biophysical responses: *Cisplatin*, a classical alkylating agent, damages the DNA in cancer cells. *Docetaxel* stabilizes microtubules, hindering their normal dynamic functions. *Erlotinib*, a second-generation targeted therapy drug, inhibits the Epidermal Growth Factor Receptor (EGFR) and its downstream signaling pathways in NSCLC. *Gemcitabine* terminates DNA elongation by binding to DNA chains as a faulty base. The inclusion of NSCLC cell lines of different subtypes, H1975 (adenocarcinoma) and H2170 (squamous cell carcinoma), demonstrates the practical application of biophysical screening in cells with distinct morphologies (**Fig. 3e, control**). The different expressions of EGFR in these cell lines (wildtype in H2170 and T790M mutated in H1975) allow for the study of distinct cellular responses to erlotinib. Each drug was tested across gradients of five concentrations with replicates, totaling 88 chambers including controls (see concentrations in **Supp. Fig. 6**, images in **Supp. Fig. 9**). Assay reproducibility was validated using intra-experimental replicates and experiments conducted across days.

Twenty-four hours prior to drug treatment, both H1975 and H2170 cells were trypsinized, resuspended in fresh culture medium, and seeded onto SpAD disks at densities of 1.2 × 10^5^ cells/mL for H1975 and 8 × 10⁴ cells/mL for H2170, respectively, with a working volume of 40 µL per chamber. Lyophilized drug compounds were reconstituted in DI water or DMSO according to their solubilities and diluted in PBS to prepare working stock solutions. Serial dilutions were subsequently performed to generate the desired concentration gradients, as detailed in **Supp. Fig. 6**. Final DMSO concentrations in all conditions were made sure to be below 0.1%. After cell attachment, the medium in each chamber was aspirated and replaced with drug-containing medium. The disk was then incubated for 24 hours before direct imaging with the SpAD system.

### MTT assay of drug-treated cells

To evaluate cell viability under the same drug treatment conditions, parallel experiments were conducted in commercial 96-well plates using the MTT Cell Proliferation Assay Kit (ThermoFisher Scientific, V13154). Cells were seeded at densities of 1.4 × 10³ cells/well for H1975 and 1.5 × 10⁴ cells/well for H2170 in a working volume of 200 µL per well. Drugs were administered using the same concentration gradients described above. At the indicated time points (24 or 96 hours post-treatment), MTT labeling and detection were performed following the manufacturer’s instructions. Between the rapid solubilization protocol using DMSO and the more complete protocol using SDS-HCl, the latter was selected. Plates were incubated at 37 °C for 18 hours after MTT addition to ensure complete solubilization of formazan crystals. Absorbance was measured at 570 nm using a plate reader. The results, averaged over four biological replicates, are presented in **Supp. Fig. 7**.

### Training-free QPI segmentation

A training-free segmentation pipeline was developed in MATLAB 2024a to robustly delineate individual cells from quantitative phase imaging (QPI) data across diverse cell types and morphological states. The algorithm integrates multiple image contrasts and segmentation techniques in a three-stage process to ensure high accuracy without requiring machine learning or manual annotation.

In the first stage, three complementary image contrasts—quantitative phase image (QPI), and the x- and y-direction differential phase gradients (DPGx, DPGy)—are utilized to generate binary masks. A simple thresholding operation is first applied to the QPI to identify the cell body and perinuclear regions, which exhibit higher phase values due to increased optical thickness. To detect thin cellular structures such as lamellipodia and filopodia—typically characterized by lower phase signals—the DPGx and DPGy images are leveraged, as they enhance contrast at boundaries and in textured regions. Masks derived from both QPI and DPG contrasts are combined to delineate the total cell-occupied area from the background in the full-field image.

In the second stage, individual cells are segmented by applying a watershed algorithm to a mask highlighting local peaks in the QPI signal, facilitating separation of closely adjacent cells. In the third stage, each preliminary cell mask is refined using an active contour model applied to the composite image gradient (ImGr), computed from the DPG contrasts. This ImGr image enhances cell boundaries with high spatial precision, allowing the active contour algorithm to converge accurately on true cellular outlines. Implementation details and source code are available at https://github.com/dsonsiu/Training-free-QPI-segmentation.

For the CRISPR screening experiment, in which the focus is on segmenting multicellular colonies rather than individual cells, the second stage was replaced with a narrow-bridge cutting algorithm, optimized for separating spatially adjacent but distinct colonies.

### Nonlinear Biophysical Trend Detection

To systematically identify nonlinear cellular responses to drug treatment, we defined and targeted three characteristic trend archetypes: monotonic shifts, hormetic responses, and sigmoid. Each archetype was detected using a dedicated algorithmic approach. Monotonic trends were assessed by computing Spearman’s rank correlation coefficient between single-cell feature values and drug concentration. Features exhibiting an absolute Spearman’s ρ > 0.1 were classified as exhibiting monotonic behavior.

For detecting hormetic and sigmoidal responses, we implemented a custom sliding window analysis inspired by the DE-SWAN framework ^35^. In this approach, Cliff’s delta was calculated between sequential concentration intervals across the gradient, generating a differential response trajectory for each feature. Hormesis was identified by a trajectory that changed direction across the concentration range (i.e., a zero-crossing pattern), indicative of biphasic behavior. Sigmoid responses were characterized by a single prominent peak in the differential trajectory, flanked by regions of near-zero effect, representing a sudden, threshold-like transition. Multiple empirically tuned thresholds were applied to robustly classify features matching each trend pattern. Chambers exhibiting atypical profiles were filtered out using mean cosine distance metrics and confirmed by visual inspection (**Supp. Fig. 10**).

### Cell culture and generation of cell lines

HEK293T cells were maintained with DMEM containing 10% FBS and 1x penicillin-streptomycin. A549-ACE2 and Vero E6-TMPRSS2 (Referred to as A549 and Vero E6 cells respectively for simplicity) cells were cultured with DMEM medium (with 10% FBS and 1x penicillin-streptomycin) supplemented with 0.5 µg ml^-1^ puromycin (ant-pr-1, InvivoGen) or 1 mg ml^-1^ G418 (ant-gn-1, InvivoGen), respectively. ACE2-, AP2M1-, FCHO2-, or CAB39 knockout cells were generated following a previously published method ^26^. All cells were incubated at 37 °C with 5% CO2.

### SpAD CRISPR Screening

Specific gene-knockout recipient cells (A549 or Vero E6) were mixed with HEK293T cells that had been transfected 24 hours earlier with SARS-CoV-2 spike-expressing plasmid (pCAG-spike-GFP11-P2A-mCherry vector) at a 1:10 ratio. These mixtures were seeded onto the SpAD disk at a final concentration of 6 × 10⁴ cells/mL and incubated at 37 °C with 5% CO₂ for 24 hours to allow cell attachment and syncytia formation.

Biological duplicates were prepared to ensure reproducibility. Following incubation, the SpAD disk was imaged first using the SpAD system and subsequently using a commercial fluorescence microscope (Nikon Ti2) under the 10X objective lens. Given the limited throughput of commercial imaging and the high consistency of cell states observed between SpAD and microscope images, fluorescence imaging was performed on only one duplicate per condition.

### Co-registering SpAD and fluorescence images

A custom image co-registration program based on MATLAB was written to co-register the images across microscopic modalities semi-automatically. Coarse overlapping was first performed manually, with 5-10 user-defined markers on both images. Next, a fine co-registration was performed automatically cell by cell. A square of 60 um margin around the cell was cropped from both images, preprocessed with a 3×3 kernel range filter and co-registered with the intensity-based image registration in MATLAB. All image registrations were optimized as rigid transforms.

### Machine Learning-Based Syncytia Classification

To distinguish syncytia from unfused colonies, machine learning (ML) models were independently developed for the A549 and Vero E6 cell lines. For each model, colonies were pooled from 11 chambers—including gene knockout (KO) conditions with matched fluorescence imaging. A stratified random split was applied to reserve half of the colonies for independent testing, ensuring class balance. The remaining colonies (A549: 3,695 unfused and 842 syncytia; Vero E6: 2,043 unfused and 373 syncytia) were used for training using the full InMorph feature set, encompassing morphological, textural, and biophysical descriptors.

Each classifier was trained using an ensemble of 30 decision trees boosted via AdaBoost, implemented in MATLAB’s Classification Learner. To assess generalization and minimize overfitting, 10-fold cross-validation was performed, yielding validation accuracies of 85.76% for A549 and 87.67% for Vero E6. Following validation, the models were evaluated on the held-out test sets and on duplicate KO chambers not subjected to fluorescence imaging. As shown in **Fig. 4e**, the ML-based classification results aligned closely with fluorescence-labeled ground truth, supporting model reliability.

### Identification of Knockout Genes Using Generalized Additive Models (GAMs)

To infer gene knockout identities from biophysical phenotypes, unfused colonies were aggregated from all KO chambers and biological replicates. Syncytia were digitally excluded using the trained ML classifiers to ensure a purified population of unfused, edited cells. Generalized Additive Models (GAMs) with Boosted Trees were selected for their capacity to model nonlinear relationships while retaining interpretability. All models were trained without score transformation and with a uniform prior; default hyperparameters were used otherwise.

Data from both A549 and Vero E6 cell lines were pooled to increase statistical power, with cell type included as a categorical predictor to account for inter-line differences. Each GAM was trained to detect colonies corresponding to a specific gene knockout. 10-fold cross-validation was performed to ensure robustness. For each colony, the trained GAMs generated a confidence score representing the likelihood of association with a given knockout. Colonies receiving consistently low scores (score < 0) across all GAMs were classified as likely derived from unedited controls. Model reliability and feature relevance were assessed via partial dependence plots, enabling biological interpretation of the learned decision boundaries.

## Notes

### Competing Interest Statement

The authors have declared no competing interest.

## References

1. Ramezani, M. et al. A genome-wide atlas of human cell morphology. Nature methods 22, 621–633 (2025).

2. Chandrasekaran, S.N. et al. Three million images and morphological profiles of cells treated with matched chemical and genetic perturbations. Nature Methods 21, 1114– 1121 (2024).

3. Lee, K.C., Guck, J., Goda, K. & Tsia, K.K. Toward deep biophysical cytometry: prospects and challenges. Trends in Biotechnology 39, 1249–1262 (2021).

4. Polanco, E.R. et al. Multiparametric quantitative phase imaging for real-time, single cell, drug screening in breast cancer. Communications Biology 5, 794 (2022).

5. Ma, H. et al. An Omni-Mesoscope for multiscale high-throughput quantitative phase imaging of cellular dynamics and high-content molecular characterization. Science Advances 10, eadq5009 (2024).

6. Cimini, B.A. et al. Optimizing the Cell Painting assay for image-based profiling. Nature protocols 18, 1981–2013 (2023).

7. Lee, K.C. et al. Multi-ATOM: Ultrahigh-throughput single-cell quantitative phase imaging with subcellular resolution. Journal of biophotonics 12, e201800479 (2019).

8. Lee, K.C. et al. Quantitative phase imaging flow cytometry for ultra-large-scale single- cell biophysical phenotyping. Cytometry Part A 95, 510–520 (2019).

9. Siu, D.M. et al. Deep-learning-assisted biophysical imaging cytometry at massive throughput delineates cell population heterogeneity. Lab on a Chip 20, 3696–3708 (2020).

10. Mansoury, M., Hamed, M., Karmustaji, R., Al Hannan, F. & Safrany, S.T. The edge effect: A global problem. The trouble with culturing cells in 96-well plates. Biochemistry and biophysics reports 26, 100987 (2021).

11. Stirling, D.R. et al. CellProfiler 4: improvements in speed, utility and usability. BMC bioinformatics 22, 433 (2021).

12. Zhang, Z. et al. Morphological profiling by high-throughput single-cell biophysical fractometry. Communications biology 6, 449 (2023).

13. Calabrese, E.J. & Mattson, M.P. How does hormesis impact biology, toxicology, and medicine? NPJ aging and mechanisms of disease 3, 13 (2017).

14. Tolis, C., Peters, G., Ferreira, C., Pinedo, H. & Giaccone, G. Cell cycle disturbances and apoptosis induced by topotecan and gemcitabine on human lung cancer cell lines. European journal of cancer 35, 796–807 (1999).

15. Hartigan, J.A. & Hartigan, P.M. The dip test of unimodality. The annals of Statistics, 70–84 (1985).

16. Pace, E. et al. Effects of gemcitabine on cell proliferation and apoptosis in non-small-cell lung cancer (NSCLC) cell lines. Cancer chemotherapy and pharmacology 46, 467– 476 (2000).

17. Wattanawongdon, W. et al. Establishment and characterization of gemcitabine-resistant human cholangiocarcinoma cell lines with multidrug resistance and enhanced invasiveness. International journal of oncology 47, 398–410 (2015).

18. Datta, A. et al. Cytoskeletal dynamics in epithelial-mesenchymal transition: insights into therapeutic targets for cancer metastasis. Cancers 13, 1882 (2021).

19. Labitigan, R.L.D. et al. Mapping variation in the morphological landscape of human cells with optical pooled CRISPRi screening. eLife 13 (2024).

20. Feldman, D. et al. Optical pooled screens in human cells. Cell 179, 787–799. e717 (2019).

21. Yan, X. et al. High-content imaging-based pooled CRISPR screens in mammalian cells. Journal of Cell Biology 220, e202008158 (2021).

22. Jiao, Y. et al. Discovering metabolic disease gene interactions by correlated effects on cellular morphology. Molecular Metabolism 24, 108–119 (2019).

23. Bussani, R. et al. Persistence of viral RNA, pneumocyte syncytia and thrombosis are hallmarks of advanced COVID-19 pathology. EBioMedicine 61 (2020).

24. Buchrieser, J. et al. Syncytia formation by SARS-CoV-2-infected cells. The EMBO journal 39, e106267 (2020).

25. Sanders, D.W. et al. SARS-CoV-2 requires cholesterol for viral entry and pathological syncytia formation. elife 10, e65962 (2021).

26. Chan, C.W. et al. High-throughput screening of genetic and cellular drivers of syncytium formation induced by the spike protein of SARS-CoV-2. Nature Biomedical Engineering 8, 291–309 (2024).

27. Seal, S. et al. Cell Painting: a decade of discovery and innovation in cellular imaging. Nature methods 22, 254–268 (2025).

28. Liu, X., Shi, L., Zhao, Z., Shu, J. & Min, W. VIBRANT: spectral profiling for single-cell drug responses. Nature methods 21, 501–511 (2024).

29. Arevalo, J. et al. Evaluating batch correction methods for image-based cell profiling. Nature Communications 15, 6516 (2024).

30. Datlinger, P. et al. Ultra-high-throughput single-cell RNA sequencing and perturbation screening with combinatorial fluidic indexing. Nature methods 18, 635–642 (2021).

31. Eckert, S. et al. Decrypting the molecular basis of cellular drug phenotypes by dose-resolved expression proteomics. Nature Biotechnology 43, 406–415 (2025).

32. Park, H.S., Ceballos, S., Eldridge, W.J. & Wax, A. Invited Article: Digital refocusing in quantitative phase imaging for flowing red blood cells. APL photonics 3 (2018).

33. Siu, D.M. et al. Optofluidic imaging meets deep learning: from merging to emerging. Lab on a Chip 23, 1011–1033 (2023).

34. Wu, W. et al. High-throughput single-cell density measurements enable dynamic profiling of immune cell and drug response from patient samples. Nature Biomedical Engineering, 1–10 (2025).

35. Lehallier, B. et al. Undulating changes in human plasma proteome profiles across the lifespan. Nature medicine 25, 1843–1850 (2019).

