## Supplementary Information for "On-The-Fly Live-Cell Intrinsic Morphological Drug and Genetic Screens by Gigapixel-per-second Spinning Arrayed Disk Imaging"

**Supplementary Note 1. Force modelling of adhered cells under spinning**

**Supplementary Table 1. Full list of InMorph features.**

**Supplementary Figure 1. SpAD optical imaging setup.**

**Supplementary Figure 2. SpAD real-time monitoring.**

**Supplementary Figure 3. Detail dimensions of SpAD disk chamber**

**Supplementary Figure 4. Focus drift.**

**Supplementary Figure 5. Comparison between SpAD and High-content screening.**

**Supplementary Figure 6. Drug concentrations.**

**Supplementary Figure 7. Drug MTT Assay Results.**

**Supplementary Figure 8. Cell Segmentation algorithm.**

**Supplementary Figure 9. Image gallery of drugged cells.**

**Supplementary Figure 10. Outlier wells removal.**

**Supplementary Figure 11. Estimated Cell Counts in SpAD Drug Experiment.**

**Supplementary Figure 12. Populations of flattened and unflattened cells**

**Supplementary Figure 13. Full ROC-AUC bar chart**

**Supplementary Figure 14. GAM replicates**

**Supplementary Note 1. Force modelling of adhered cells under spinning**

To model the forces experienced by the adhered cells inside a spinning SpAD, we consider **a rotating frame of reference** (the frame co-moving with the chamber and the medium of SpAD). In total, four relevant forces act upon the cell:

**A. Centrifugal Force**

$$\vec{F}_{cent} = m_{cell}\omega^2\vec{r}$$

where $m_{cell}$ is cell mass, $\omega$ is angular velocity, $\vec{r}$ is a position vector relative to the rotational axis.

**B. Buoyant Force**

$$\vec{F}_{buoy} = -m_{medium,displaced}\omega^2\vec{r}$$

where the mass of the fluid displaced by the cell is $m_{medium,displaced} = \rho_{medium}V_{cell}$. In this case, $\rho_{medium}$ is the density of the fluid medium, and $V_{cell}$ is the volume of the cell.

Since the surrounding culture medium is also set in a rotating motion, a counteracting buoyant force is exerted onto the cell by the surrounding medium. The **buoyant force** acts in the opposite direction of the centrifugal force.

**C. Coriolis Force**

$$\vec{F}_{cori} = -2m_{cell}(\vec{\omega} \times \vec{v}_{rel})$$

where $\vec{v}_{rel}$ is the relative velocity of the cell relative to the rotating reference frame.

It is relevant only **if the cell is moving with respect to the rotating frame** (i.e., if the cell moves relative to the fluid/chamber).

But in our scenario, the adhered cell is regarded stationary or "very-slow-varying" relative to the rotating chamber and medium, so ,$\vec{v}_{rel} \approx 0$ and Coriolis force $\vec{F}_{cori} \approx 0$.

**D. Euler Force**

$$\vec{F}_{Euler} = -m_{cell}(\frac{d\vec{\omega}}{dt} \times \vec{r})$$

Euler force emerges when the rotational speed changes. Thus, during the course of the experiment (i.e. constant rotational velocity), $\frac{d\vec{\omega}}{dt} \approx 0$ and Euler force $\vec{F}_{Euler} \approx 0$.

*Note: If the axis of rotation is not aligned with gravity, gravity acts independently, but is usually negligible compared to centrifugal force at high rpm.*

Consider a practical scenario shown below, standard cell culture medium (e.g., DMEM, RPMI) supplemented with 10% FBS has a density of $\sim 1.00 - 1.03\ g/cm^3$ [1] whereas mammalian cell normally has density of $\sim 1.05\ g/cm^3$ [2]. Hence, the density difference is

$\Delta\rho = \rho_{cell} - \rho_{medium} = 0.02\ g/cm^3$

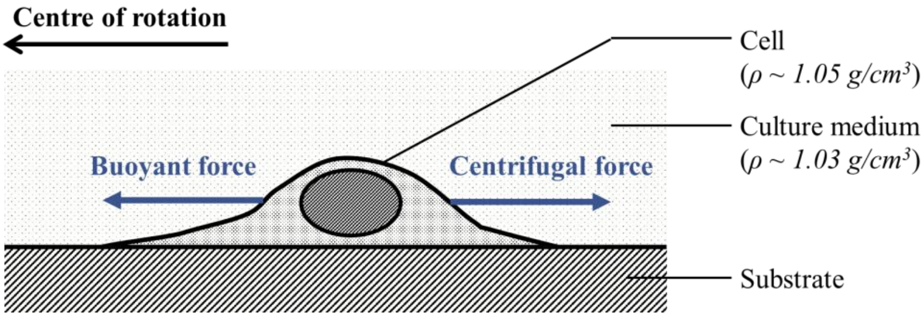


Further consider the following parameters in the SpAD scenario,

> **Radius:** $r = 50mm$
> **Spin speed:** $f = 1000\ rpm \Rightarrow \omega = 2\pi f = 104.72\ rad/s$
> **Cell volume:** $V_{cell} \approx 2000\ \mu m^3$ (~15 µm in diameter).

Therefore, the magnitude of the net force is $F_{net} = 2.19 \times 10^{-11}\ N$

For comparison, the **weight of a typical cell** under gravity ($F_g = mg = 2.06 \times 10^{-11}$ N). So the net force from spinning is comparable to the cell's weight on Earth—but only because of the density difference.

**This is also orders of magnitude smaller the typical *whole* cell adhesion force: $10^{-8} - 10^{-7}$ N [3, 4]

### Supplementary Table 1. Full list of InMorph features.

| | |
|---|---|
| 1 | CellMorpho area |
| 2 | CellMorpho perimeter |
| 3 | CellMorpho circularity |
| 4 | CellMorpho eccentricity |
| 5 | CellMorpho orientation |
| 6 | CellMorpho major axis length |
| 7 | CellMorpho minor axis length |
| 8 | CellMorpho aspect ratio |
| 9 | CellMorpho max feret diameter |
| 10 | CellMorpho min feret diameter |
| 11 | CellMorpho feret diameters ratio |
| 12 | CellMorpho convex area |
| 13 | CellMorpho equivalent diameter |
| 14 | CellMorpho extent |
| 15 | CellMorpho solidity |
| 16 | CellMorpho compactness |
| 17 | CellMorpho mean radius |
| 18 | CellMorpho max radius |
| 19 | CellMorpho min radius |
| 20 | CellMorpho Zernike_0_0 |
| 21 | CellMorpho Zernike_1_1 |
| 22 | CellMorpho Zernike_2_0 |
| 23 | CellMorpho Zernike_2_2 |
| 24 | CellMorpho Zernike_3_1 |
| 25 | CellMorpho Zernike_3_3 |
| 26 | CellMorpho Zernike_4_0 |
| 27 | CellMorpho Zernike_4_2 |
| 28 | CellMorpho Zernike_4_4 |
| 29 | CellMorpho Zernike_5_1 |
| 30 | CellMorpho Zernike_5_3 |
| 31 | CellMorpho Zernike_5_5 |
| 32 | CellMorpho Zernike_6_0 |
| 33 | CellMorpho Zernike_6_2 |
| 34 | CellMorpho Zernike_6_4 |
| 35 | CellMorpho Zernike_6_6 |
| 36 | CellMorpho Zernike_7_1 |
| 37 | CellMorpho Zernike_7_3 |
| 38 | CellMorpho Zernike_7_5 |
| 39 | CellMorpho Zernike_7_7 |
| 40 | CellMorpho Zernike_8_0 |
| 41 | CellMorpho Zernike_8_2 |
| 42 | CellMorpho Zernike_8_4 |
| 43 | CellMorpho Zernike_8_6 |
| 44 | CellMorpho Zernike_8_8 |
| 45 | CellMorpho Zernike_9_1 |
| 46 | CellMorpho Zernike_9_3 |
| 47 | CellMorpho Zernike_9_5 |
| 48 | CellMorpho Zernike_9_7 |
| 49 | CellMorpho Zernike_9_9 |
| 50 | CellMorpho HuMoment1 |
| 51 | CellMorpho HuMoment2 |
| 52 | CellMorpho HuMoment3 |
| 53 | CellMorpho HuMoment4 |
| 54 | CellMorpho HuMoment5 |
| 55 | CellMorpho HuMoment6 |
| 56 | CellMorpho HuMoment7 |
| 57 | CellMorpho InertiaTensor11 |
| 58 | CellMorpho InertiaTensor22 |
| 59 | CellMorpho InertiaTensor12 |
| 60 | CellMorpho InerTensEigenVal 1 |
| 61 | CellMorpho InerTensEigenVal 2 |
| 62 | Number of seeds |
| 63 | Total Area of seeds |
| 64 | Seeds Area Mean |
| 65 | Seeds Area Variance |
| 66 | Seeds Area Skewness |
| 67 | Seeds Area Kurtosis |
| 68 | Seeds Solidity Mean |
| 69 | Seeds Solidty Variance |
| 70 | Seeds Solidty Skewness |
| 71 | Seeds Solidty Kurtosis |
| 72 | Seeds Eccentricity Mean |
| 73 | Seeds Eccentricity Variance |
| 74 | Seeds Eccentricity Skewness |
| 75 | Seeds Eccentricity Kurtosis |
| 76 | Seeds Perimeter Mean |
| 77 | Seeds Perimeter Variance |
| 78 | Seeds Perimeter Skewness |
| 79 | Seeds Perimeter Kurtosis |
| 80 | Seeds Circularity Mean |
| 81 | Seeds Circularity Variance |
| 82 | Seeds Circularity Skewness |
| 83 | Seeds Circularity Kurtosis |
| 84 | Seed-seed Distance Mean |
| 85 | Seed-seed Distance Variance |
| 86 | Seed-seed Distance Skewness |
| 87 | Seed-seed Distance Kurtosis |
| 88 | Seeds-cellcen Distance Mean |
| 89 | Seeds-cellcen Distance Variance |
| 90 | Seeds-cellcen Distance Skewness |
| 91 | Seeds-cellcen Distance Kurtosis |
| 92 | Seed-cell Area Ratio |
| 93 | SDMap mean |
| 94 | SDMap var |
| 95 | SDMap skewness |
| 96 | SDMap kurtosis |
| 97 | SDMap range |
| 98 | SDMap max |
| 99 | SDMap min |
| 100 | SDMap centroid shift |
| 101 | SDMap radial distribution |
| 102 | SDMap Entr1um mean |
| 103 | SDMap Entr1um var |
| 104 | SDMap Entr1um skewness |
| 105 | SDMap Entr1um kurtosis |
| 106 | SDMap Entr1um range |
| 107 | SDMap Entr1um max |
| 108 | SDMap Entr1um min |
| 109 | SDMap Entr1um centroid shift |
| 110 | SDMap Entr1um radial distribution |
| 111 | SDMap Entr2um mean |
| 112 | SDMap Entr2um var |
| 113 | SDMap Entr2um skewness |
| 114 | SDMap Entr2um kurtosis |
| 115 | SDMap Entr2um range |
| 116 | SDMap Entr2um max |
| 117 | SDMap Entr2um min |
| 118 | SDMap Entr2um centroid shift |
| 119 | SDMap Entr2um radial distribution |
| 120 | SDMap Entr5um mean |
| 121 | SDMap Entr5um var |
| 122 | SDMap Entr5um skewness |
| 123 | SDMap Entr5um kurtosis |
| 124 | SDMap Entr5um range |
| 125 | SDMap Entr5um max |
| 126 | SDMap Entr5um min |
| 127 | SDMap Entr5um centroid shift |
| 128 | SDMap Entr5um radial distribution |
| 129 | SDMap Entr10um mean |
| 130 | SDMap Entr10um var |
| 131 | SDMap Entr10um skewness |
| 132 | SDMap Entr10um kurtosis |
| 133 | SDMap Entr10um range |
| 134 | SDMap Entr10um max |
| 135 | SDMap Entr10um min |
| 136 | SDMap Entr10um centroid shift |
| 137 | SDMap Entr10um radial distribution |
| 138 | SDMap std1um mean |
| 139 | SDMap std1um var |
| 140 | SDMap std1um skewness |
| 141 | SDMap std1um kurtosis |
| 142 | SDMap std1um range |
| 143 | SDMap std1um max |
| 144 | SDMap std1um min |
| 145 | SDMap std1um centroid shift |
| 146 | SDMap std1um radial distribution |
| 147 | SDMap std2um mean |
| 148 | SDMap std2um var |
| 149 | SDMap std2um skewness |
| 150 | SDMap std2um kurtosis |
| 151 | SDMap std2um range |
| 152 | SDMap std2um max |
| 153 | SDMap std2um min |
| 154 | SDMap std2um centroid shift |
| 155 | SDMap std2um radial distribution |
| 156 | SDMap std5um mean |
| 157 | SDMap std5um var |
| 158 | SDMap std5um skewness |
| 159 | SDMap std5um kurtosis |
| 160 | SDMap std5um range |
| 161 | SDMap std5um max |
| 162 | SDMap std5um min |
| 163 | SDMap std5um centroid shift |
| 164 | SDMap std5um radial distribution |
| 165 | SDMap std10um mean |
| 166 | SDMap std10um var |
| 167 | SDMap std10um skewness |
| 168 | SDMap std10um kurtosis |
| 169 | SDMap std10um range |
| 170 | SDMap std10um max |
| 171 | SDMap std10um min |
| 172 | SDMap std10um centroid shift |
| 173 | SDMap std10um radial distribution |
| 174 | SDMap range1um mean |
| 175 | SDMap range1um var |
| 176 | SDMap range1um skewness |
| 177 | SDMap range1um kurtosis |
| 178 | SDMap range1um range |
| 179 | SDMap range1um max |
| 180 | SDMap range1um min |
| 181 | SDMap range1um centroid shift |
| 182 | SDMap range1um radial distribution |
| 183 | SDMap range2um mean |
| 184 | SDMap range2um var |
| 185 | SDMap range2um skewness |
| 186 | SDMap range2um kurtosis |
| 187 | SDMap range2um range |
| 188 | SDMap range2um max |
| 189 | SDMap range2um min |
| 190 | SDMap range2um centroid shift |
| 191 | SDMap range2um radial distribution |
| 192 | SDMap range5um mean |
| 193 | SDMap range5um var |
| 194 | SDMap range5um skewness |
| 195 | SDMap range5um kurtosis |
| 196 | SDMap range5um range |
| 197 | SDMap range5um max |
| 198 | SDMap range5um min |
| 199 | SDMap range5um centroid shift |
| 200 | SDMap range5um radial distribution |
| 201 | SDMap range10um mean |
| 202 | SDMap range10um var |
| 203 | SDMap range10um skewness |
| 204 | SDMap range10um kurtosis |
| 205 | SDMap range10um range |
| 206 | SDMap range10um max |
| 207 | SDMap range10um min |
| 208 | SDMap range10um centroid shift |
| 209 | SDMap range10um radial distribution |
| 210 | SDMap Fiber mean |
| 211 | SDMap Fiber var |
| 212 | SDMap Fiber skewness |
| 213 | SDMap Fiber kurtosis |
| 214 | SDMap Fiber 50Percen |
| 215 | SDMap Fiber 75Percen |
| 216 | SDMap Fiber centroid shift |
| 217 | SDMap Fiber radial distribution |
| 218 | DMMap mean |
| 219 | DMMap var |
| 220 | DMMap skewness |
| 221 | DMMap kurtosis |
| 222 | DMMap range |
| 223 | DMMap max |
| 224 | DMMap min |
| 225 | DMMap centroid shift |
| 226 | DMMap radial distribution |
| 227 | DMMap Entr1um mean |
| 228 | DMMap Entr1um var |
| 229 | DMMap Entr1um skewness |
| 230 | DMMap Entr1um kurtosis |
| 231 | DMMap Entr1um range |
| 232 | DMMap Entr1um max |
| 233 | DMMap Entr1um min |
| 234 | DMMap Entr1um centroid shift |
| 235 | DMMap Entr1um radial distribution |
| 236 | DMMap Entr2um mean |
| 237 | DMMap Entr2um var |
| 238 | DMMap Entr2um skewness |
| 239 | DMMap Entr2um kurtosis |
| 240 | DMMap Entr2um range |
| 241 | DMMap Entr2um max |
| 242 | DMMap Entr2um min |
| 243 | DMMap Entr2um centroid shift |
| 244 | DMMap Entr2um radial distribution |
| 245 | DMMap Entr5um mean |
| 246 | DMMap Entr5um var |
| 247 | DMMap Entr5um skewness |
| 248 | DMMap Entr5um kurtosis |
| 249 | DMMap Entr5um range |
| 250 | DMMap Entr5um max |
| 251 | DMMap Entr5um min |
| 252 | DMMap Entr5um centroid shift |
| 253 | DMMap Entr5um radial distribution |
| 254 | DMMap Entr10um mean |
| 255 | DMMap Entr10um var |
| 256 | DMMap Entr10um skewness |
| 257 | DMMap Entr10um kurtosis |
| 258 | DMMap Entr10um range |
| 259 | DMMap Entr10um max |
| 260 | DMMap Entr10um min |
| 261 | DMMap Entr10um centroid shift |
| 262 | DMMap Entr10um radial distribution |
| 263 | DMMap std1um mean |
| 264 | DMMap std1um var |
| 265 | DMMap std1um skewness |
| 266 | DMMap std1um kurtosis |
| 267 | DMMap std1um range |
| 268 | DMMap std1um max |
| 269 | DMMap std1um min |
| 270 | DMMap std1um centroid shift |
| 271 | DMMap std1um radial distribution |
| 272 | DMMap std2um mean |
| 273 | DMMap std2um var |
| 274 | DMMap std2um skewness |
| 275 | DMMap std2um kurtosis |
| 276 | DMMap std2um range |
| 277 | DMMap std2um max |
| 278 | DMMap std2um min |
| 279 | DMMap std2um centroid shift |
| 280 | DMMap std2um radial distribution |
| 281 | DMMap std5um mean |
| 282 | DMMap std5um var |
| 283 | DMMap std5um skewness |
| 284 | DMMap std5um kurtosis |
| 285 | DMMap std5um range |
| 286 | DMMap std5um max |
| 287 | DMMap std5um min |
| 288 | DMMap std5um centroid shift |
| 289 | DMMap std5um radial distribution |
| 290 | DMMap std10um mean |
| 291 | DMMap std10um var |
| 292 | DMMap std10um skewness |
| 293 | DMMap std10um kurtosis |
| 294 | DMMap std10um range |
| 295 | DMMap std10um max |
| 296 | DMMap std10um min |
| 297 | DMMap std10um centroid shift |
| 298 | DMMap std10um radial distribution |
| 299 | DMMap range1um mean |
| 300 | DMMap range1um var |
| 301 | DMMap range1um skewness |
| 302 | DMMap range1um kurtosis |
| 303 | DMMap range1um range |
| 304 | DMMap range1um max |
| 305 | DMMap range1um min |
| 306 | DMMap range1um centroid shift |
| 307 | DMMap range1um radial distribution |
| 308 | DMMap range2um mean |
| 309 | DMMap range2um var |
| 310 | DMMap range2um skewness |
| 311 | DMMap range2um kurtosis |
| 312 | DMMap range2um range |
| 313 | DMMap range2um max |
| 314 | DMMap range2um min |
| 315 | DMMap range2um centroid shift |
| 316 | DMMap range2um radial distribution |
| 317 | DMMap range5um mean |
| 318 | DMMap range5um var |
| 319 | DMMap range5um skewness |
| 320 | DMMap range5um kurtosis |
| 321 | DMMap range5um range |
| 322 | DMMap range5um max |
| 323 | DMMap range5um min |
| 324 | DMMap range5um centroid shift |
| 325 | DMMap range5um radial distribution |
| 326 | DMMap range10um mean |
| 327 | DMMap range10um var |
| 328 | DMMap range10um skewness |
| 329 | DMMap range10um kurtosis |
| 330 | DMMap range10um range |
| 331 | DMMap range10um max |
| 332 | DMMap range10um min |
| 333 | DMMap range10um centroid shift |
| 334 | DMMap range10um radial distribution |
| 335 | DMMap Fiber mean |
| 336 | DMMap Fiber var |
| 337 | DMMap Fiber skewness |
| 338 | DMMap Fiber kurtosis |
| 339 | DMMap Fiber 50Percen |
| 340 | DMMap Fiber 75Percen |
| 341 | DMMap Fiber centroid shift |
| 342 | DMMap Fiber radial distribution |
| 343 | DMRadMap mean |
| 344 | DMRadMap variance |
| 345 | DMRadMap skewness |
| 346 | DMAngFreq variance |
| 347 | DMAngFreq kurtosis |
| 348 | BFQPI Correlation |
| 349 | BFQPI Overlap coef |
| 350 | BFQPI Manders coef (SDM) |
| 351 | BFQPI Manders coef (QPI) |
| 352 | ALS mean |
| 353 | ALS variance |
| 354 | ALS skewness |
| 355 | ALS kurtosis |
| 356 | ALS range |
| 357 | ALS max |
| 358 | ALS 0.05 intensity drop |
| 359 | ALS 0.1 intensity drop |
| 360 | ALS first min height |
| 361 | ALS max peak difference |
| 362 | ALS slope peak drop |
| 363 | ALS max slope |
| 364 | Fractal dimension |
| 365 | Fractal dim within window |
| 366 | Fractal window width |
| 367 | Fractal fit error 1 |
| 368 | Fractal fit error 2 |
| 369 | Mean distance of nearest ten |
| 370 | Distance to nearest |
| 371 | Number in range |
| 372 | Adjacency length |
| 373 | Proportion of adjacent perimeter |
| 374 | Adjacenciness |
| 375 | Number of adjacent cells |
| 376 | Dry mass |
| 377 | Cell flatness |

**Feature categories legend**

| | |
|---|---|
| Bulk | Dry mass global |
| Cortex bulk | Dry mass local |
| Scatterer global | Colocalization |
| Scatterer local | Cell-cell interaction |

We categorized the extracted features into eight major types (see legend) to comprehensively characterize single-cell phenotypes. Following established morphological profiling approaches [5], basic shape descriptors were used to define overall cell morphology and are labeled as *bulk features*. To specifically assess the cell cortex—where most organelles reside and which exhibits elevated phase values—we defined a separate group of *cortex bulk features*.

*Scatterer features* (annotated as **SDMap**, for scatterer density map) were derived from brightfield (BF) images. These capture dark puncta representing intracellular materials that absorb or scatter light. *Dry mass features* (annotated as **DMMap**, for dry mass map) were computed from quantitative phase images (QPI), leveraging the known relationship between optical phase shift and dry mass density [6].

*Global features* describe overall pixel intensity distributions within each cell, while *local features* were extracted using multi-scale spatial filters to highlight subcellular structures at varying length scales [7]. *Colocalization features* quantify the spatial overlap between scatterer and dry mass signals, offering insight into intracellular organization.

*Angular light scattering (ALS)* and *fractal features* reflect whole-cell scattering behaviors, which can only be computed when both amplitude (BF) and phase (QPI) information are available [8]. Finally, *cell–cell interaction features* assess the proximity and arrangement of neighboring cells.

Together, this panel of features enables a high-dimensional, label-free phenotypic profiling of each individual cell.

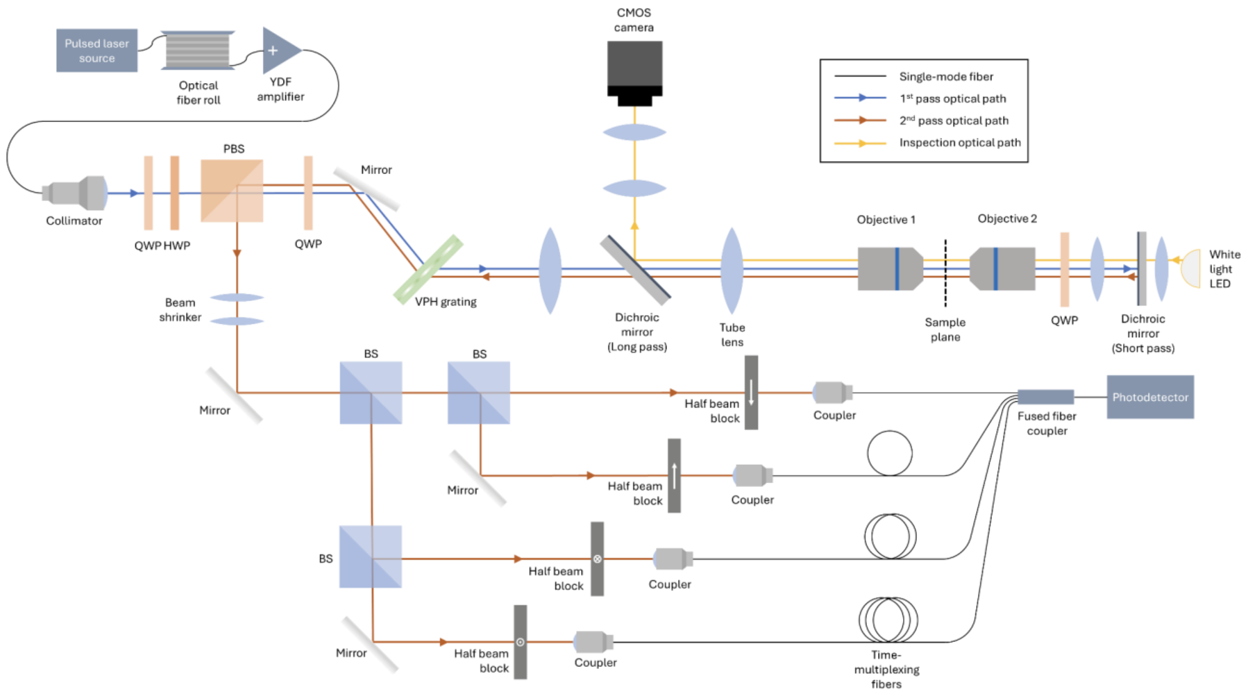


**Supplementary Figure 1. SpAD optical imaging setup.**

The optical imaging setup of SpAD was built based on Multi-ATOM [9]. Compared to the last reported system, minor modifications and improvements were incorporated to better accommodate SpAD. Firstly, the collimator, relay lenses and objectives combination was selected to engineer the beam diameter and the effective NA to 0.56, to optimize FOV and resolution without compromising the QPI accuracy. Secondly, a lens was added between objective 2 and the short-pass dichroic mirror to maintain the 4-f in this double-pass configuration, which minimizes aberrations. QWP: Quarter-wave plate; HWP: Half-wave plate; PBS: Polarization beamsplitter; VPH grating: Volume phase holographic grating; BS: Beamsplitter.

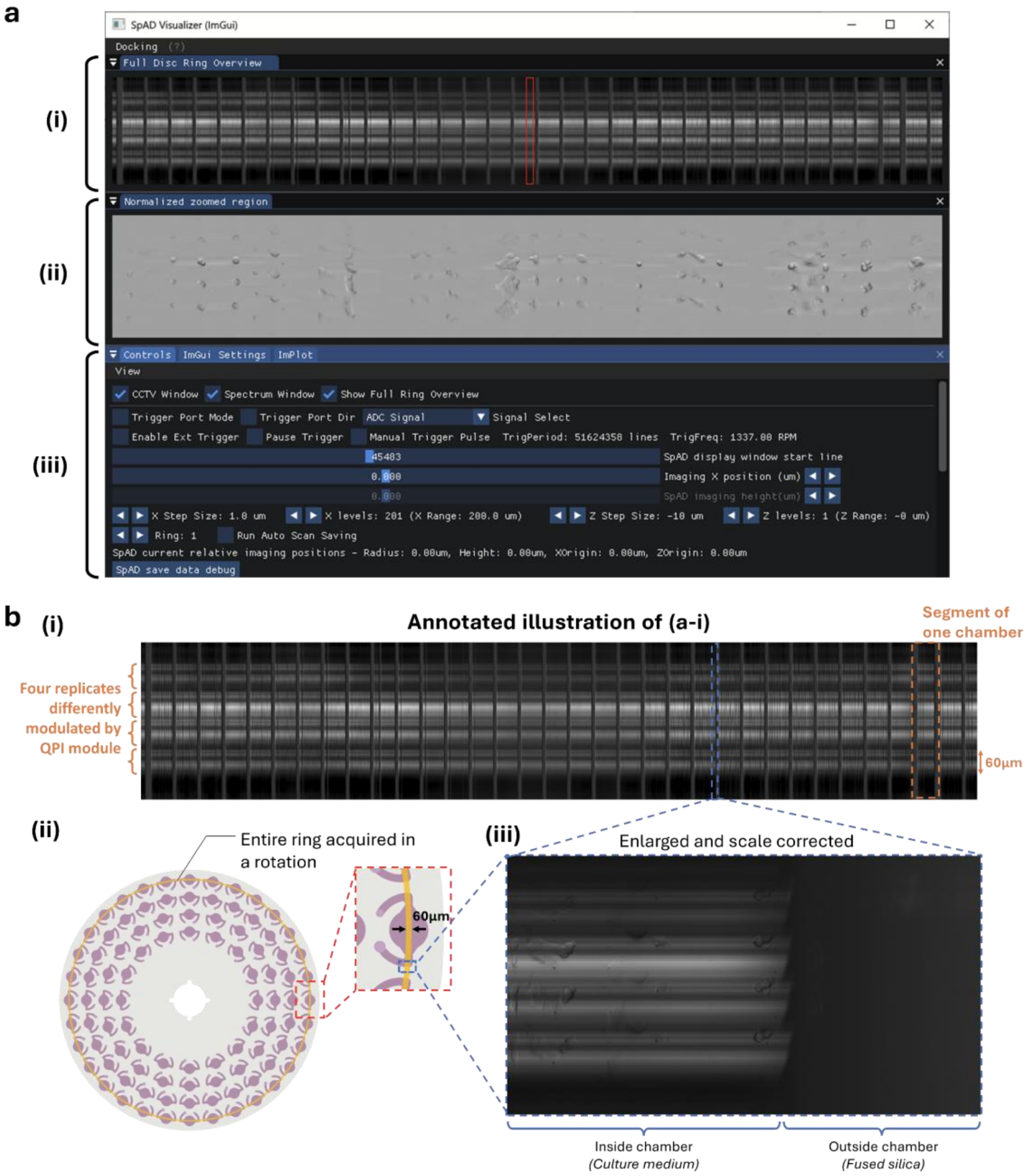


**Supplementary Figure 2. SpAD real-time monitoring interface.**

To facilitate high quality image acquisition, we develop **(a)** a custom user interface for real-time monitoring and automatic data acquisition. **(a-i)** The top overview window shows a straightened, horizontally compressed image of the entire ring. In this encoded image, the horizontal axis defines the angular position of the SpAD. The vertical axis encodes four modulated replicates of the same imaging region. **(a-ii)** Zoomed-in view of the selected region (red box) from *(a-i)*, shown in the correct aspect ratio. The image is intensity-normalized to correct for spectral variation and improve visibility. Users can scroll horizontally to inspect different regions of the ring in real time. *(a-i)* and *(a-ii)* update with ~20fps according to the rotation speed, which is enough for fine focus tuning. **(a-iii)** Control panel enabling adjustments to SpAD translation, visualization options, and acquisition parameters. **(b-i)** Annotated overview of the ring image shown in *(a-i)*, where the vertical axis represents four modulated replicates of the same imaging region. Horizontal bright and dark bands arise from spectral intensity fluctuations of the pulsed laser. The orange dashed box marks a single chamber segment. **(b-ii)** Illustration of the ring region corresponding to the visualized overview in *(a-i)* and *(b-i)*. **(b-iii)** Enlarged view of the chamber edge within the selected segment (Blue box). Areas between chambers contain fused silica rather than culture medium. Due to refractive index mismatch, light from these regions does not couple back into the detector, resulting in dark vertical bands.

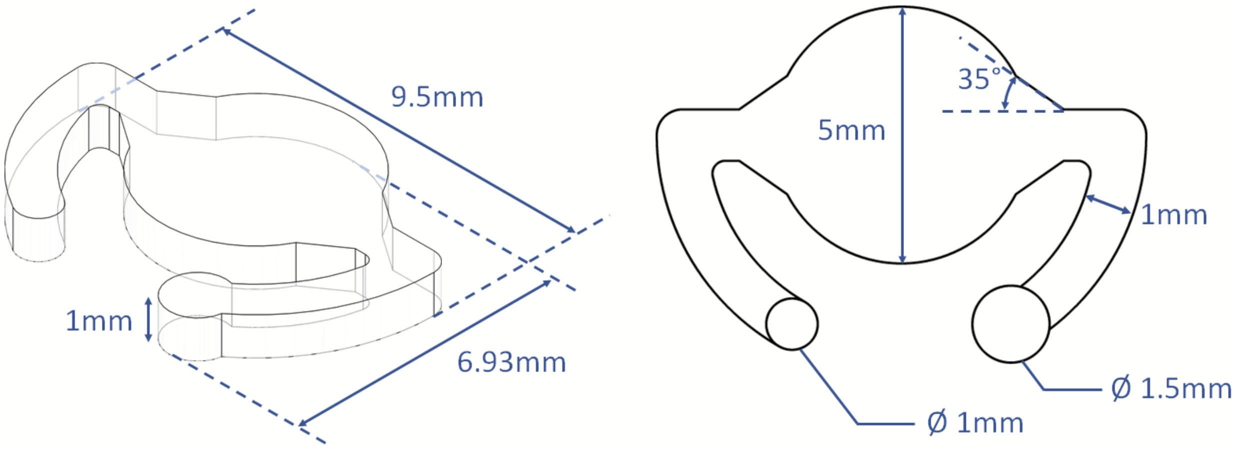


**Supplementary Figure 3. Detail dimensions of SpAD disk chamber**

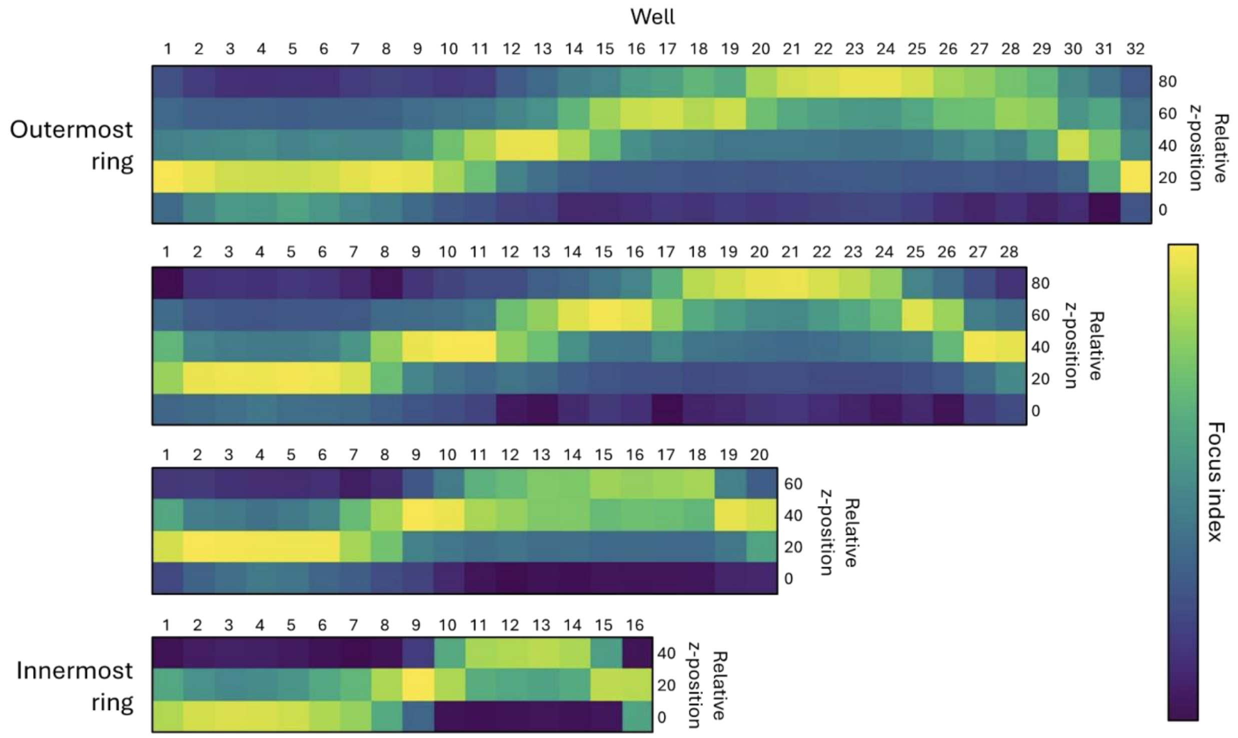


**Supplementary Figure 4. Focus drift.**

Z-stack images were acquired for each of the four rings of wells in SpAD, with step size of 20µm. To calculate the focus index, the reconstructed image of the well was first subject to *imgradient()* to obtain the image gradient magnitude map. Then, the 95th percentile of the image histogram was selected as the focus factor of the well. This calculation is based on the assumption that focused images produce the sharpest image gradient which is true in most image contrasts. The heat map of the outermost ring shows that the focus drift in the range from 20 to 80µm only for wells with centres located 55mm from the rotational centre. The wobbling is ~0.0625°.

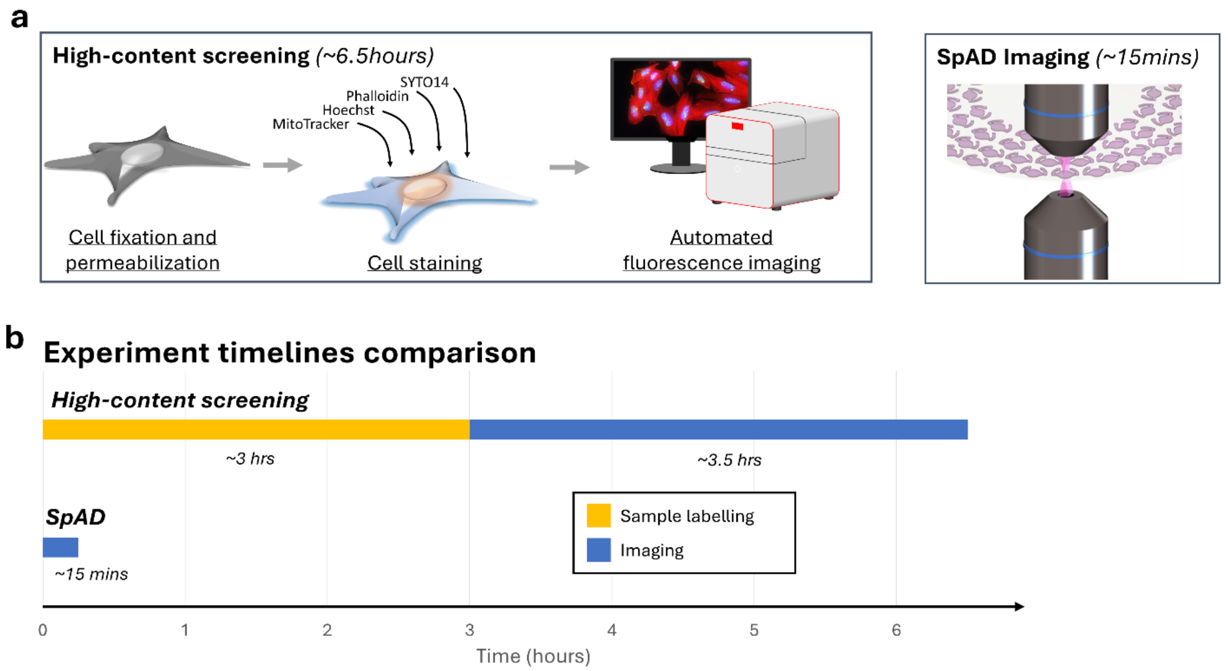


**Supplementary Figure 5. Comparison between SpAD and High-content screening.**

**(a)** Schematic overview of the workflow for conventional high-content screening versus SpAD-based screening. **(b)** Comparison of experimental timelines between the two approaches. SpAD significantly reduces total assay time by eliminating sample labelling steps and maximizing imaging throughput. Timing and procedures for conventional screening are referenced from [10, 11].

| Concentration gradient | Cisplatin | | Docetaxel | | Erlotinib | | Gemcitabine | |
|---|---|---|---|---|---|---|---|---|
| | C1 | 1.75 µM | D1 | 0.0625 nM | E1 | 0.167 µM | G1 | 0.8 nM |
| | C2 | 3.5 µM | D2 | 0.5 nM | E2 | 0.5 µM | G2 | 4 nM |
| | C3 | 7 µM | D3 | 4 nM | E3 | 1.5 µM | G3 | 20 nM |
| | C4 | 14 µM | D4 | 32 nM | E4 | 4.5 µM | G4 | 100 nM |
| | C5 | 28 µM | D5 | 256 nM | E5 | 13.5 µM | G5 | 500 nM |

**Supplementary Figure 6. Drug concentrations.**

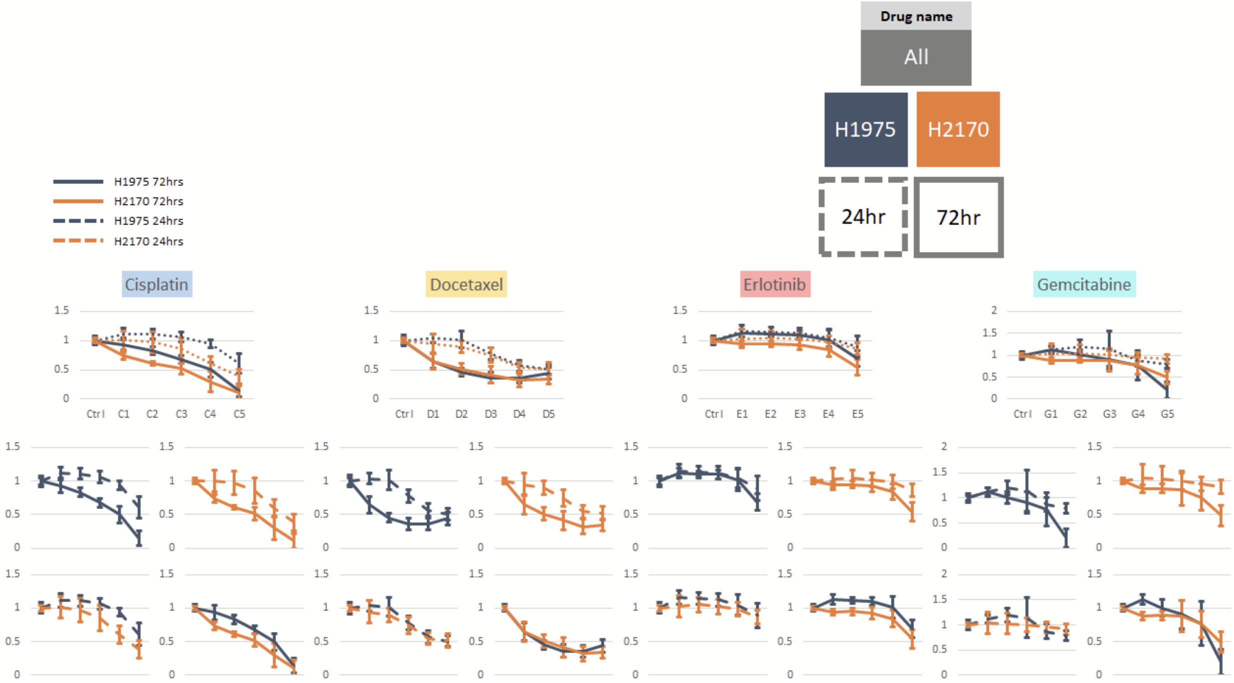


**Supplementary Figure 7. Drug MTT Assay Results.**

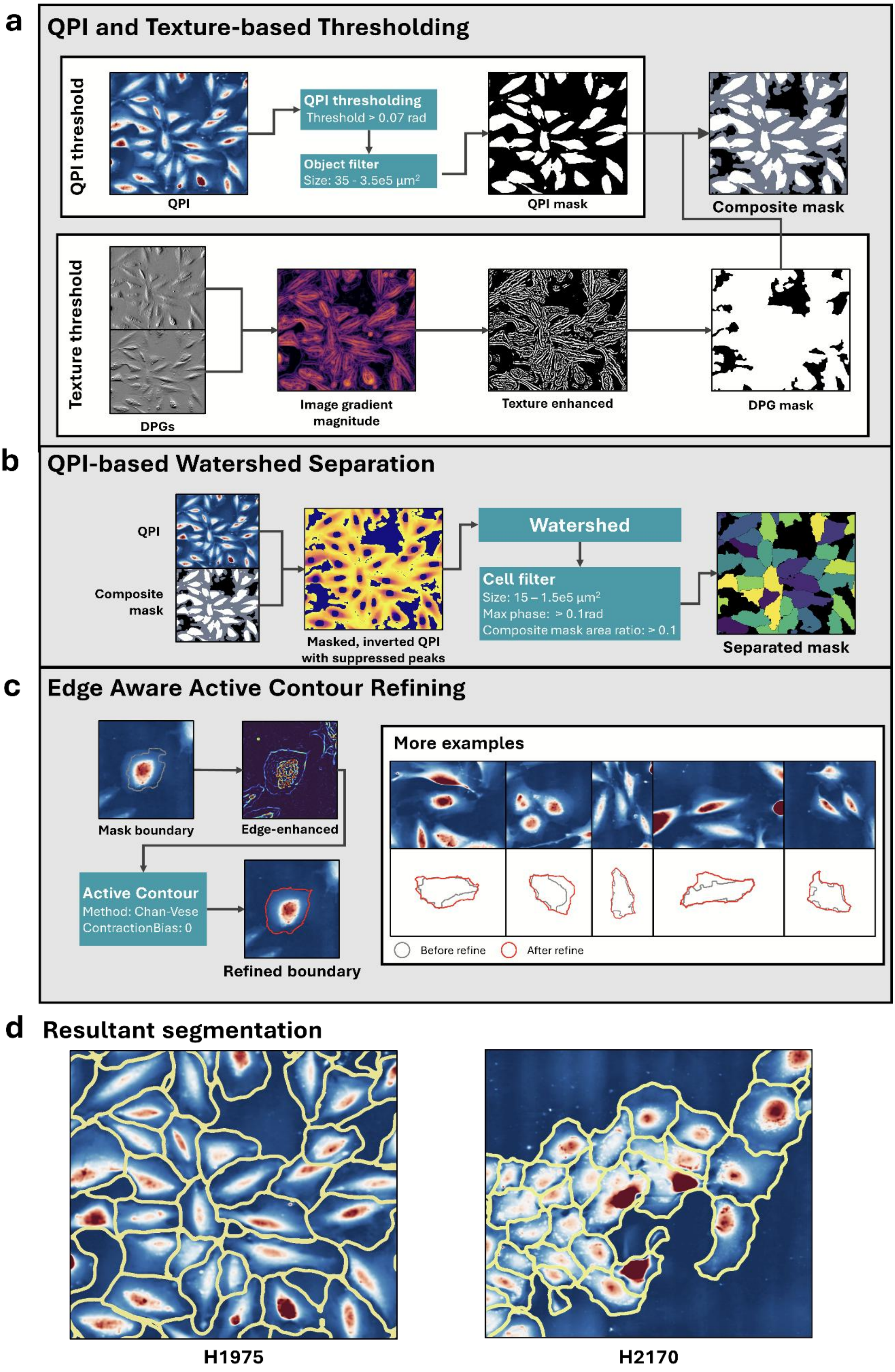
a
QPI and Texture-based Thresholding
QPI threshold
QPI
QPI thresholding
Threshold > 0.07 rad
Object filter
Size: 35 - 3.5e5 μm²
QPI mask
Composite mask
Texture threshold
DPGs
Image gradient magnitude
Texture enhanced
DPG mask
b
QPI-based Watershed Separation
QPI
Composite mask
Masked, inverted QPI with suppressed peaks
Watershed
Cell filter
Size: 15 – 1.5e5 μm²
Max phase: > 0.1rad
Composite mask area ratio: > 0.1
Separated mask
c
Edge Aware Active Contour Refining
Mask boundary
Edge-enhanced
Active Contour
Method: Chan-Vese
ContractionBias: 0
Refined boundary
More examples
Before refine
After refine
d
Resultant segmentation
H1975
H2170

**Supplementary Figure 8. Cell Segmentation algorithm.**

(a) QPI and texture-based thresholding. (b) QPI-based watershed separation. (c) Edge-aware active contour refining.

QPI provides pseudo-3D information indicative of the nucleus position and cell boundaries, while the phase gradient image emphasizes local textures that highlight intracellular organelles, distinguishing the cell body from background regions. Our QPI-based segmentation makes use of these features through segregating the process into three stages (**Fig. 3d**), each optimized for a specific task: (1) *Level-Thresholding:* In the first stage, we apply a level-thresholding method to the QPI and phase gradient images to segment cells from the background. (2) *Watershed Algorithm*: Next, we implement the watershed algorithm to separate individual cells based on local peaks appearing in the phase image, which indicate the cell body. (3) *Active Contour*: Finally, we use active contour to refine cell boundaries, ensuring that the segmentation accurately depicts cell morphology.

This integrated, multi-stage approach is inherently generalizable to all 2D cultures, as it relies on universal cellular features rather than dataset-specific training. By assigning each task to the algorithm best suited for it, our method outperforms traditional approaches that rely on a single technique while avoiding the computational and data-intensive requirements of deep learning.

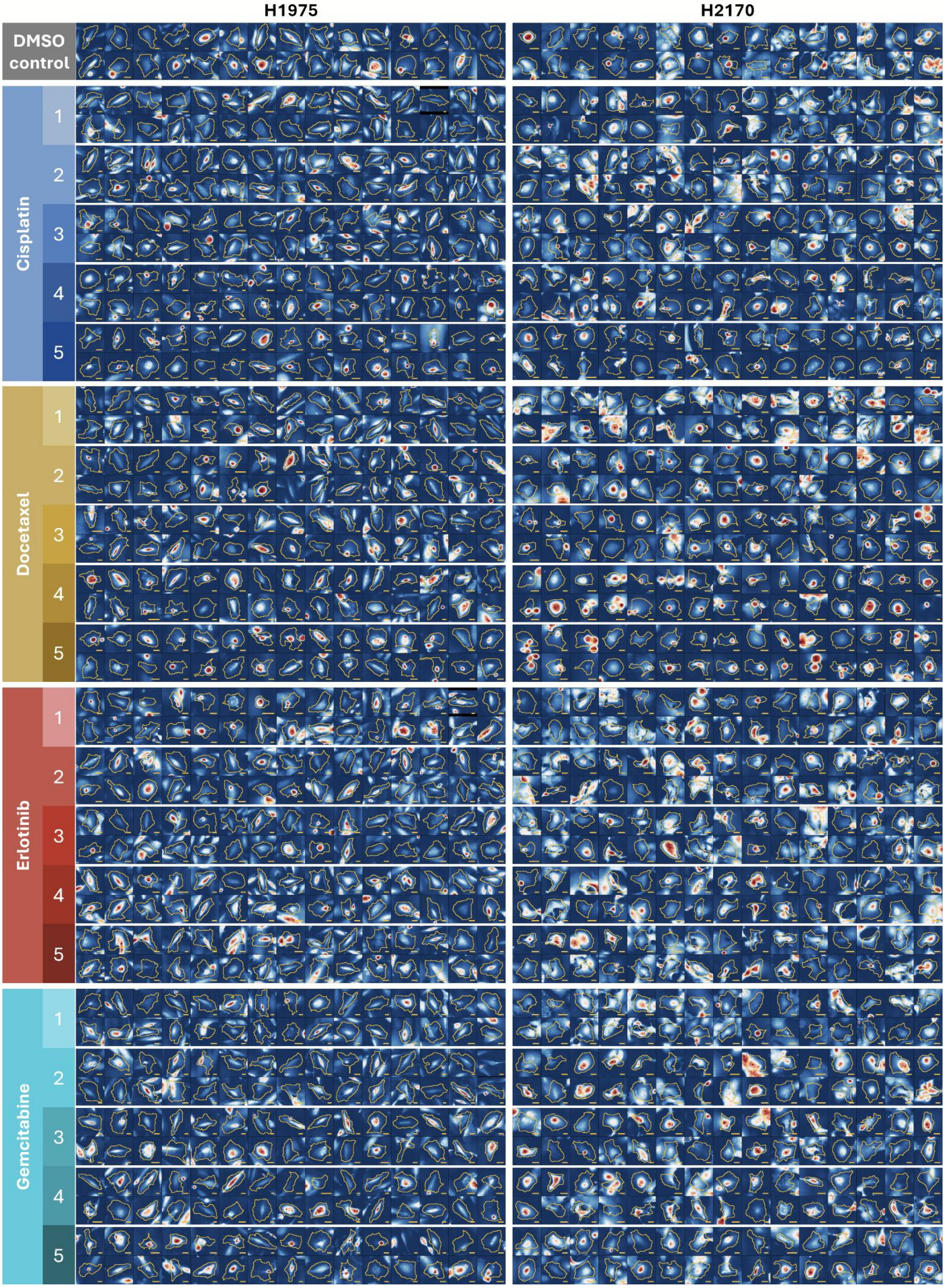


**Supplementary Figure 9. Image gallery of drugged cells.**

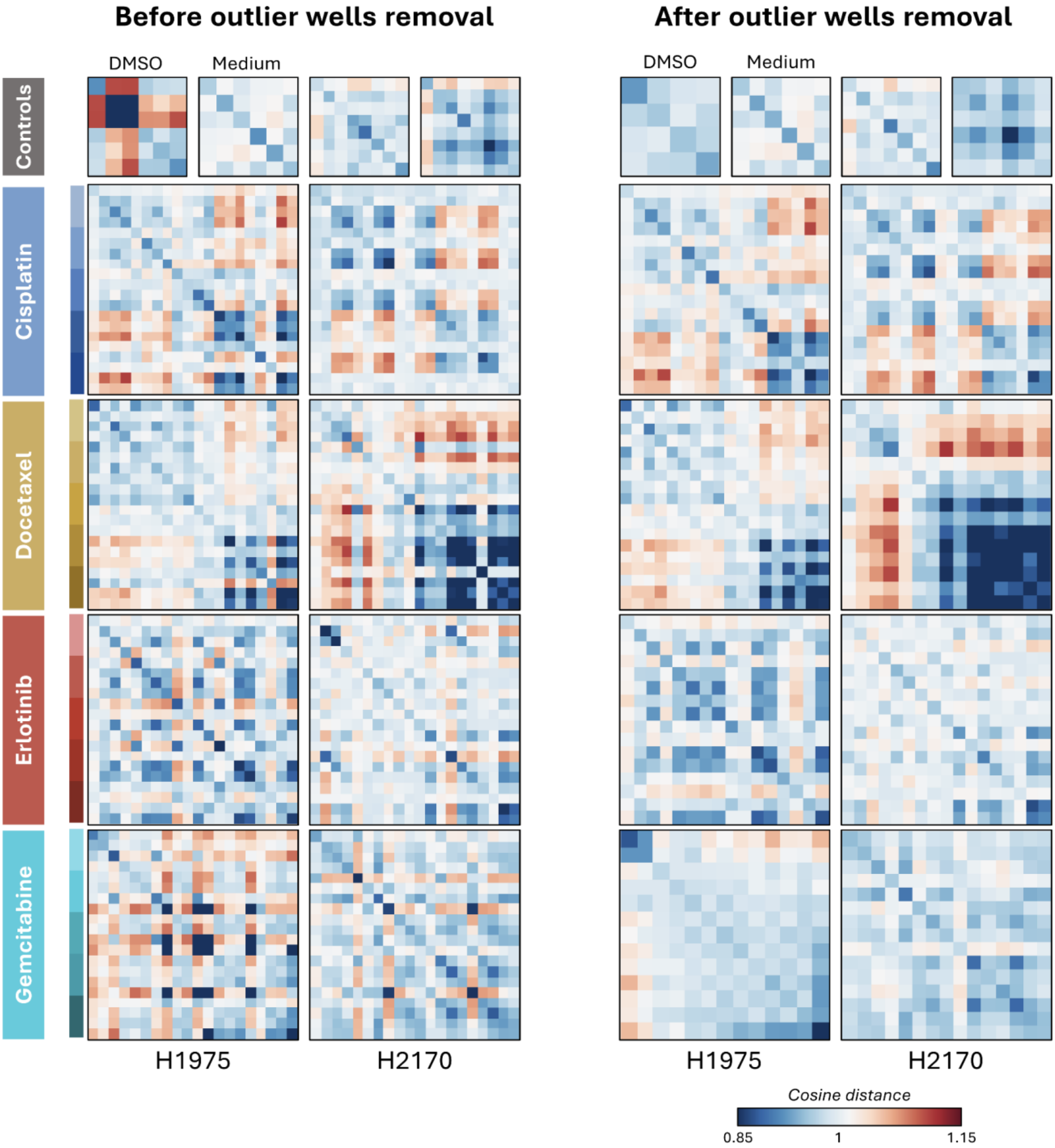


**Supplementary Figure 10. Outlier wells removal.**

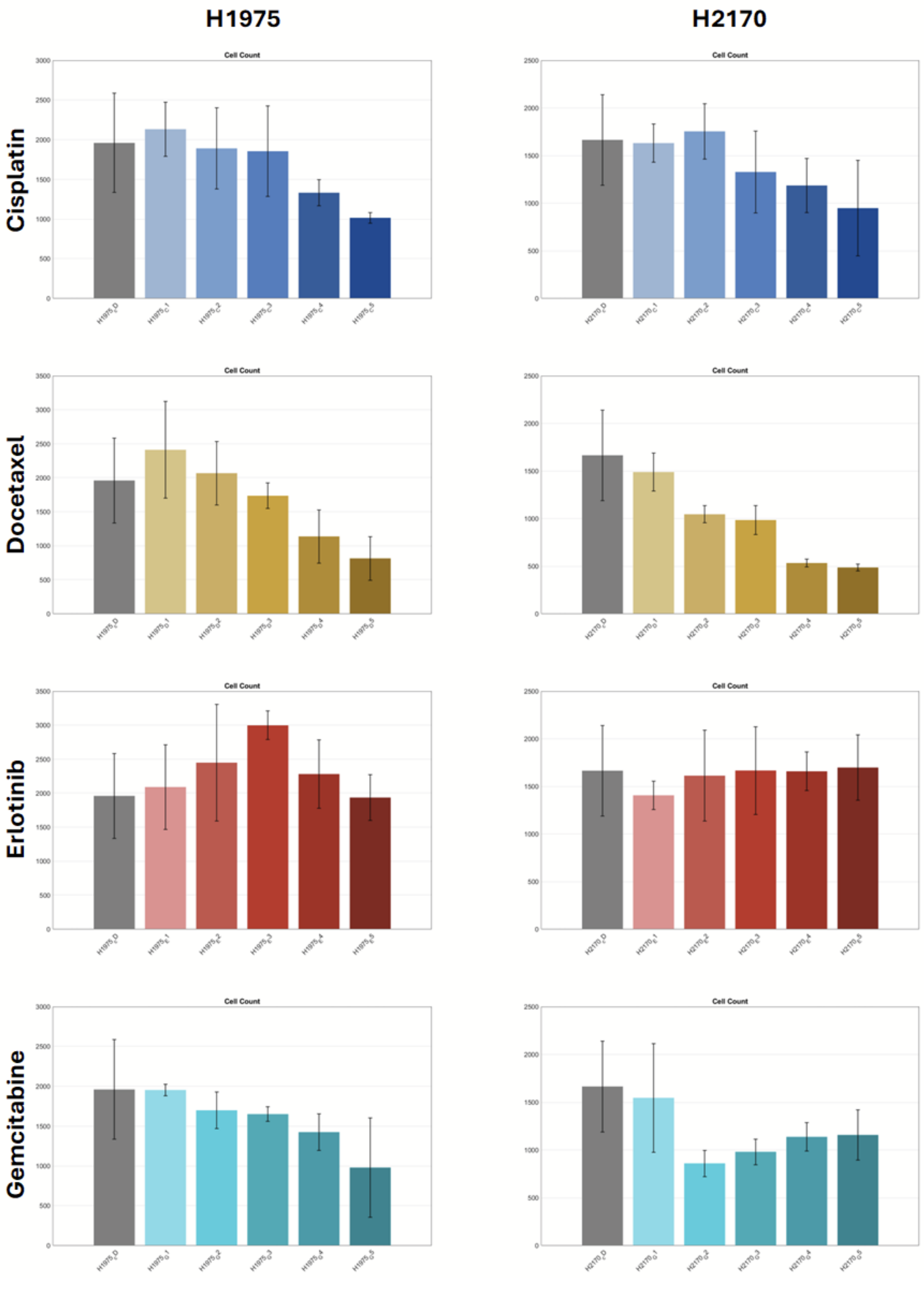


**Supplementary Figure 11. Estimated Cell Counts in SpAD Drug Experiment.**

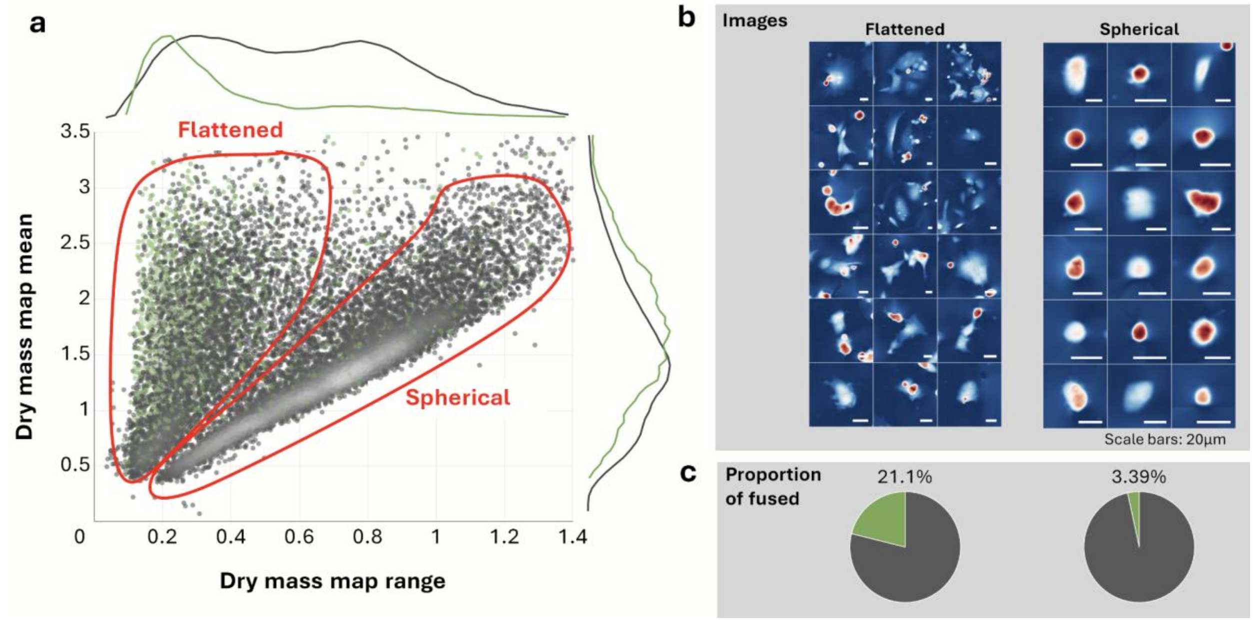


**Supplementary Figure 12. Populations of flattened and unflattened cells**

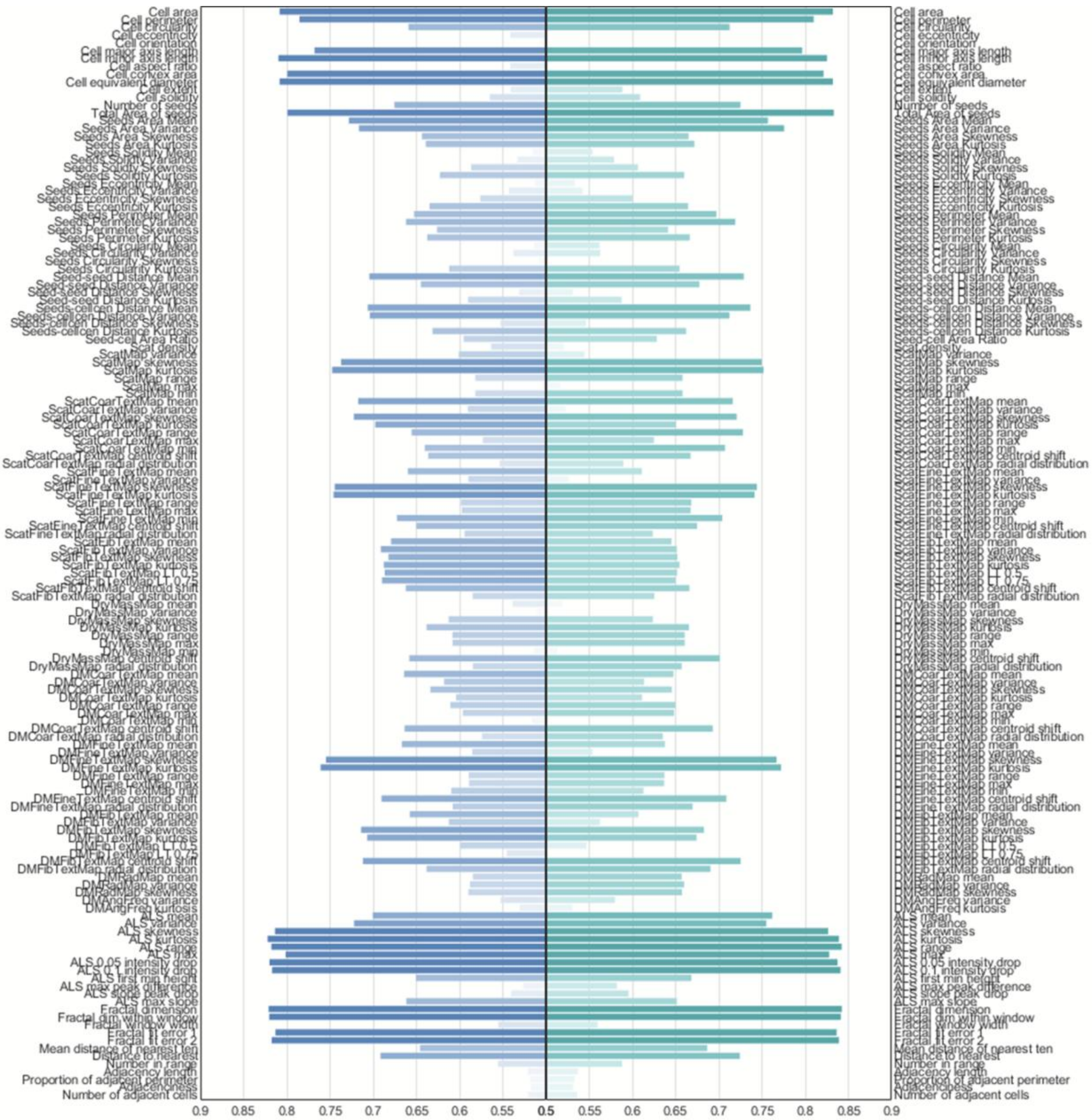


**Supplementary Figure 13. Full ROC-AUC bar chart**

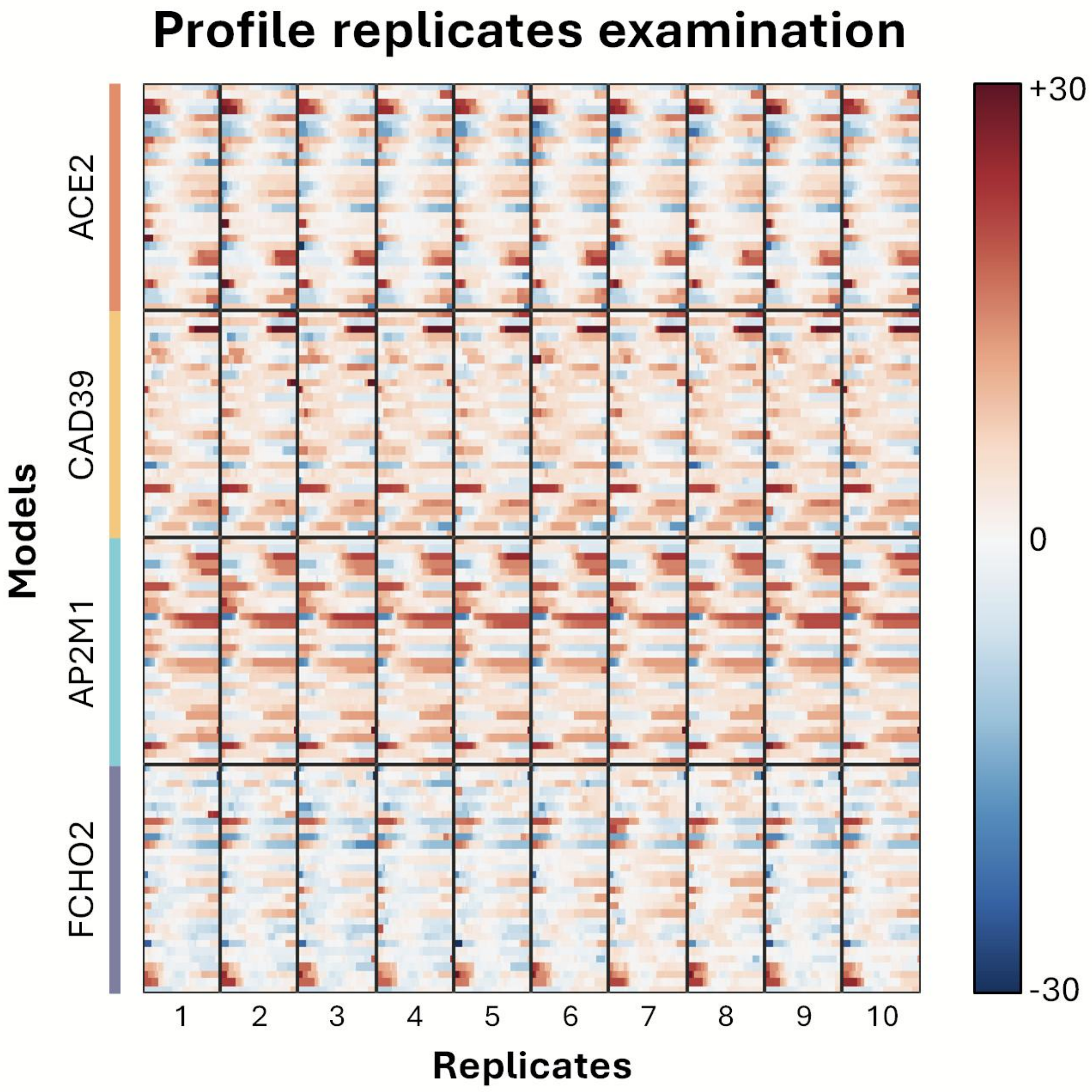


**Supplementary Figure 14. GAM replicates**

### References


1. Poon, C. Measuring the density and viscosity of culture media for optimized computational fluid dynamics analysis of in vitro devices. *Journal of the mechanical behavior of biomedical materials* **126**, 105024 (2022).
2. Bryan, A.K. et al. Measuring single cell mass, volume, and density with dual suspended microchannel resonators. *Lab on a Chip* **14**, 569–576 (2014).
3. Sztilkovics, M. et al. Single-cell adhesion force kinetics of cell populations from combined label-free optical biosensor and robotic fluidic force microscopy. *Scientific reports* **10**, 61 (2020).
4. Ungai-Salánki, R. et al. A practical review on the measurement tools for cellular adhesion force. *Advances in colloid and interface science* **269**, 309–333 (2019).
5. Stirling, D.R. et al. CellProfiler 4: improvements in speed, utility and usability. *BMC bioinformatics* **22**, 433 (2021).
6. Goswami, N., Anastasio, M.A. & Popescu, G. Quantitative phase imaging techniques for measuring scattering properties of cells and tissues: a review–part II. *Journal of biomedical optics* **29**, S22714–S22714 (2024).
7. Siu, D.M. et al. Deep-learning-assisted biophysical imaging cytometry at massive throughput delineates cell population heterogeneity. *Lab on a Chip* **20**, 3696–3708 (2020).
8. Zhang, Z. et al. Morphological profiling by high-throughput single-cell biophysical fractometry. *Communications biology* **6**, 449 (2023).
9. Lee, K.C. et al. Multi-ATOM: Ultrahigh-throughput single-cell quantitative phase imaging with subcellular resolution. *Journal of biophotonics* **12**, e201800479 (2019).
10. Bray, M.-A. et al. Cell Painting, a high-content image-based assay for morphological profiling using multiplexed fluorescent dyes. *Nature protocols* **11**, 1757–1774 (2016).
11. Cimini, B.A. et al. Optimizing the Cell Painting assay for image-based profiling. *Nature protocols* **18**, 1981–2013 (2023).
